# Genome-wide targets identification by CRISPRi-Omics for high-titer production of free fatty acids in *Escherichia coli*

**DOI:** 10.1101/2020.03.03.974378

**Authors:** Lixia Fang, Jie Fan, Congya Wang, Yingxiu Cao, Hao Song

## Abstract

To construct a superior microbial cell factory for chemical synthesis, a major challenge is to fully exploit cell potential via identifying and engineering beneficial gene targets in the sophisticated metabolic networks. Here, we develop an approach that integrates CRISPR interference (CRISPRi) to readily modulate genes expression and omics analyses to identify potential targets in multiple cellular processes, enabling systematical discovery of beneficial chromosomal gene targets that can be engineered to optimize free fatty acids (FFAs) production in *Escherichia coli*. We identify 56 beneficial genes via synergistic CRISPRi-Omics strategy, including 46 novel targets functioning in cell structure and division, and signaling transduction that efficiently facilitate FFAs production. Upon repressing *ihfA* and overexpressing *aidB* and *tesA’* in *E. coli*, the recombinant strain LihfA-OaidB results in a FFAs titer of 21.6 g L^-1^ in fed-batch fermentation, which, to our best knowledge, is the maximum FFAs titer by the recombinant *E. coli* reported to date.

## Introduction

Microbial biosynthesis of a desired product is a sophisticated process, usually involving the coordination of multiple metabolic intermediates or reactions subject to complicated interactions^1,2^. On the one hand, each of these intermediates flows to other metabolic processes for biosynthesis of cellular components or metabolites, acting as one of the multi-branched nodes in the topology of metabolic landscapes^3^. On the other hand, each reaction in the biosynthetic pathway deeply rely on many cellular processes, such as genes transcription and translation to enable enzymes function^4-6^, or cell proliferation to ensure biomass accumulation^7^. As such, manipulation of other metabolic pathways or cellular processes might lead to an unexpectedly improvement on the biosynthesis of a desired product due to distant effects of genetic modulations or unknown regulatory interactions^8,9^. Thus, to construct a superior microbial cell factory for an enhanced product biosynthesis, it is highly desirable to genome-widely identify beneficial gene targets in the highly complicated intracellular interaction networks that can be reengineered for this purpose. As such, the interwoven linkage mechanism between product biosynthesis and other sophisticated genetic and metabolic networks can be better understood.

Free fatty acids (FFAs) are feedstocks for the manufacture of detergents, lubricants, cosmetics, and pharmaceutical ingredients^10,11^. A number of metabolic engineering approaches were developed to facilitate FFAs production in *E. coli*. Overexpression of key enzymes in the FFAs biosynthetic pathway such as acetyl-coenzyme A (CoA) carboxylase^12-14^ and fatty acyl-ACP thioesterase^12,13,15-17^, or deletion of key genes in the beta-oxidation pathway such as *fadD*^12,13,15-17^ or *fadE*^15^ gene, were studied. In particular, optimally combinatorial expression of 15 crucial genes in the FFAs biosynthetic pathway with *fadD* deletion in *E. coli* BL21 resulted in a titer of 8.6 g L^-1^ of FFAs^18^. However, the cell potential for FFAs overproduction is still not fully unleashed via modulating the FFAs directly related metabolic pathways alone^19^. Besides, the cellular rewiring mechanism and the linkage between FFAs biosynthesis and other cellular processes is still largely unknown, and it remains difficult in systematically identifying beneficial gene targets using laborious and time-consuming genome engineering methods, such as homologous recombination-based gene knockout^20^. These hurdles severely restrict the speed and scope of engineering of microbial hosts to obtain a superior overproducer.

Here, we employ a systems metabolic engineering strategy for rapid, systematical and effective identification of chromosomal gene targets that can be engineered for the enhanced production of FFAs in *E. coli*. This strategy makes use of both the CRISPR interference (CRISPRi) that readily down-regulates genes expression in a dose-dependent manner to rapidly identify beneficial knockdown targets, and the omics analyses of different strains that allows acquisition of comprehensive data related to mechanisms of cellular metabolism to dig potential targets in the genetic and metabolic networks. By way of this synergistic CRISPRi–Omics strategy, we identified 46 new beneficial gene targets that efficiently facilitate FFAs production, which involves in cell division, signaling transduction and transport, cell structure, and other seemingly irrelevant cellular processes with FFAs biosynthesis. Upon repressing *ihfA* and overexpressing *aidB* and *tesA’*, the resulting recombinant *E. coli* strain LihfA-OaidB enabled a FFAs titer of 21.6 g L^-1^ (with productivity of 0.636 g L^-1^ h^-1^) in fed-batch fermentation, which, to the best of our knowledge, is the maximally reported FFAs titer and productivity by the recombinant *E. coli* to date. Our findings not only allow for enhanced biosynthesis of the desired product, but also lead to a mechanistic understanding of the underlying linkage between product biosynthesis and multiple cellular processes.

## Results

### Construction of a non-toxic and functional CRISPRi system in *E. coli* BL21(DE3)

CRISPRi system functioned through the expression of dCas9 and customized specific sgRNA^21^ (**Supplementary Fig. 1a**). However, high-level dCas9 could cause a severe inhibition of cell growth and abnormal expression of genes in a strain-specific manner^22,23^. To determine a superior dCas9 expression level in BL21(DE3), we constructed plasmids with dCas9 being expressed under the control of the promoter P_T7_, P_T5_, P_Trc_ and P_BAD_, respectively (**Supplementary Fig. 2a**). The expression strengths of these promoters were P_T7_ > P_T5_ > P_Trc_ > P_BAD_, according to our previous study^24^. With green fluorescent protein (GFP) as a characterization signal driven by the P_T7_ promoter, fluorescence-based reporter plasmids were further constructed (**Supplementary Fig. 2a**) and transformed into BL21(DE3), forming five strains G0 (P_T7_-GFP), G1 (P_T7_-GFP-P_T7_-dCas9), G2 (P_T7_-GFP-P_T5_-dCas9), G3 (P_T7_-GFP-P_Trc_-dCas9), and G4 (P_T7_-GFP-P_BAD_-dCas9), respectively. We found that the OD_600_ and fluorescence level of the strains G1 and G2 were greatly decreased from the strain G0, while that of G3 and G4 were little affected compared with G0 (**Supplementary Fig. 1b**). This suggested that the expression of dCas9 at relatively low level, such as under the control of the P_Trc_ or P_BAD_ promoter, did not cause overt toxicity in our system.

Then, we examined the transcriptional repression capability of CRISPRi on gene expression by co-expressing a specific sgRNA. sgRNA-*gfp* was designed to target on the non-template strand of GFP at a position ∼38 nucleotides downstream of the start codon to expect a high repression efficiency^25^. The plasmid Sg-Hgfp (**Supplementary Fig. 2b**), harboring sgRNA-*gfp* under the control of the P_R_ promoter, was constructed and transformed into the strains G0-G4, forming the strains g0-g4, respectively. The results showed that fluorescence levels of the strains co-expressing dCas9 and sgRNA-*gfp* (g1-g4) were all dramatically reduced in comparison to the corresponding strain expressing only dCas9 (G1-G4) (**Supplementary Fig. 1b**). It indicated that the CRISPRi system was functional for effective transcriptional repression. In addition, the strain with dCas9 regulated by the P_Trc_ promoter obtained the highest repression efficiency (93%) (**Supplementary Fig. 1b**). Thus, we selected dCas9 controlled by the P_Trc_ promoter to construct the CRISPRi system in BL21(DE3).

### Identification of beneficial gene targets in the pathways related to FFAs metabolism using CRISPRi system

Glycerol, serving as the substrate, is catabolized into key intermediates glyceraldehyde-3P, pyruvate, acetyl-CoA, malonyl-ACP in sequence and then converted into acyl-ACP through the fatty acid biosynthetic pathway (**Fig. 1a**). However, these key intermediates also involved in other metabolic pathways, resulting in diverted carbon fluxes away from FFAs. The expression of truncated fatty acyl-ACP thioesterase TesA’ (with leader sequence deleted) could release free fatty acid intermediates from ACP in the cytosol^26,27^. Nevertheless, the synthetic FFAs are degraded naturally by the beta-oxidation process (**Fig. 1a**). To channel carbon flux towards FFAs and impede products’ degradation, we chose 108 genes in the chromosome that related with the FFAs metabolism and regulation as candidate targets for modulation by CRISPRi, and classified them into five modules (upstream carbon flux diversion, downstream carbon flux diversion, amino acid metabolism, beta-oxidation, and transcription factors) (**Fig. 1a**).

**Fig. 1.**
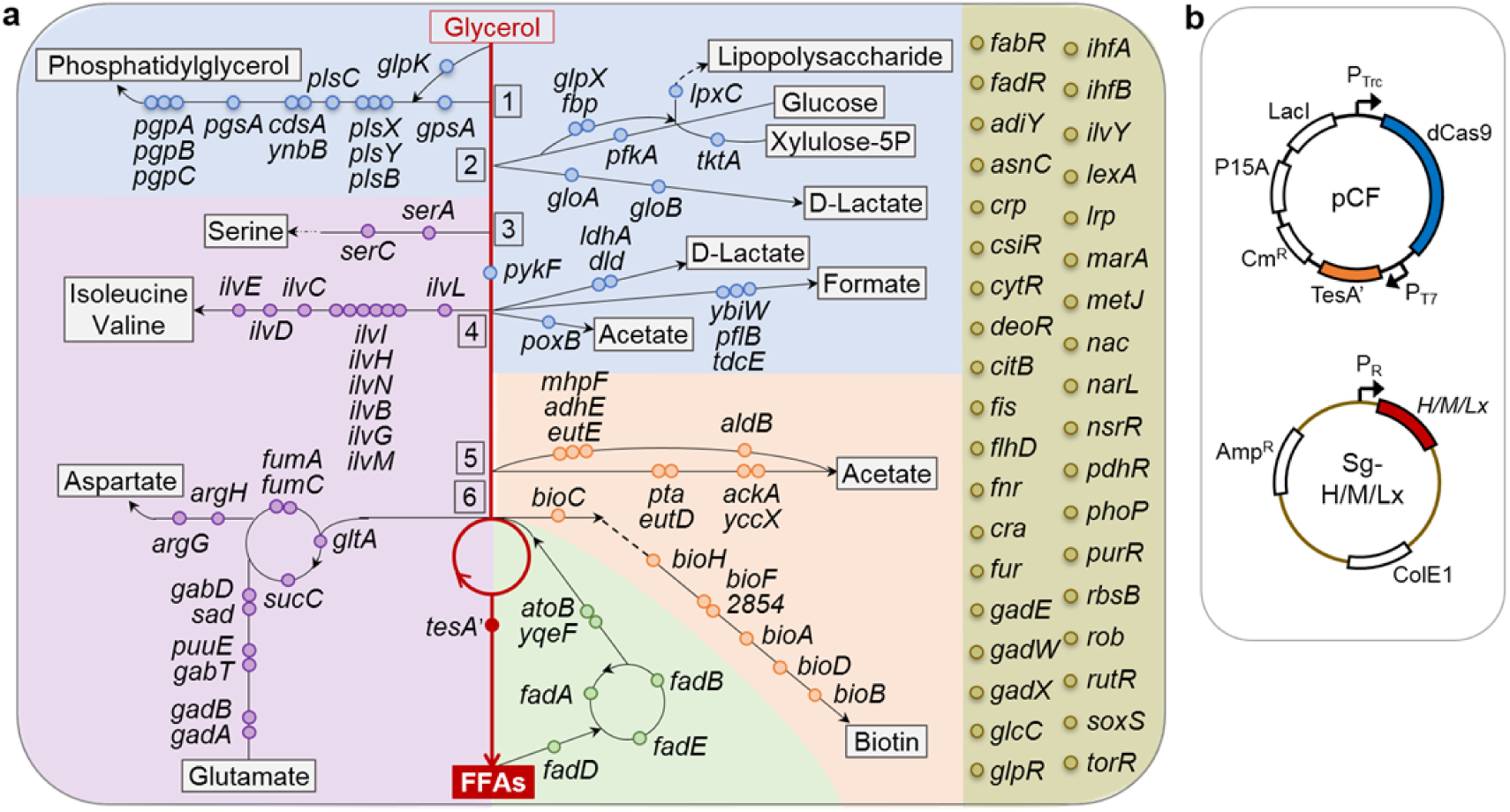
The CRISPRi approach for identification of beneficial knockdown gene targets from cellular pathways associated with the FFAs metabolism. **a** Schematic of the metabolic or regulatory pathways related to FFAs biosynthesis in *E. coli*, which were classified into five modules (upstream carbon flux diversion, downstream carbon flux diversion, amino acid metabolism, beta-oxidation and transcription factors) and represented in blue, orange, purple, green and golden background, respectively. Genes in the above five modules are shown in italics and marked with blue, orange, purple, green and golden dot, respectively. *2854* represents gene numbered ECD_02854 in BL21(DE3). The fatty acid biosynthetic pathway is shown in red line and key metabolites glycerone-P, glyceraldehyde-3P, glycerate-3P, pyruvate, acetyl-CoA and malonyl-ACP are represented as number 1 to 6, respectively. **b** Plasmid constructs for the gene expression of *tesA’, dCas9* and synthetic sgRNAs.

To facilitate large-scale target identification, we constructed a library of 108 synthetic sgRNAs that repress the expression of the 108 genes in the above five modules with high silencing efficiency^21^. We introduced plasmids harboring each of the 108 synthetic sgRNAs individually into the starting strain CF co-expressing TesA’ and dCas9 in BL21(ED3) to implement gene perturbations (**Fig. 1b**). The strain CF transformed with plasmid Sg-0 that expressed sgRNA0 without target site complementary sequence was named as the strain Control, which produced FFAs with a tier of 631 mg L^-1^. Any CRISPRi-engineered strain that could increase FFAs titer by more than 20% compared with the strain Control was sorted out. As a result, 30 beneficial targets were identified from the library and the highest production of FFAs reached 1,232 mg L^-1^, 95% higher than that of the strain Control (**Fig. 2**).

**Fig. 2.**
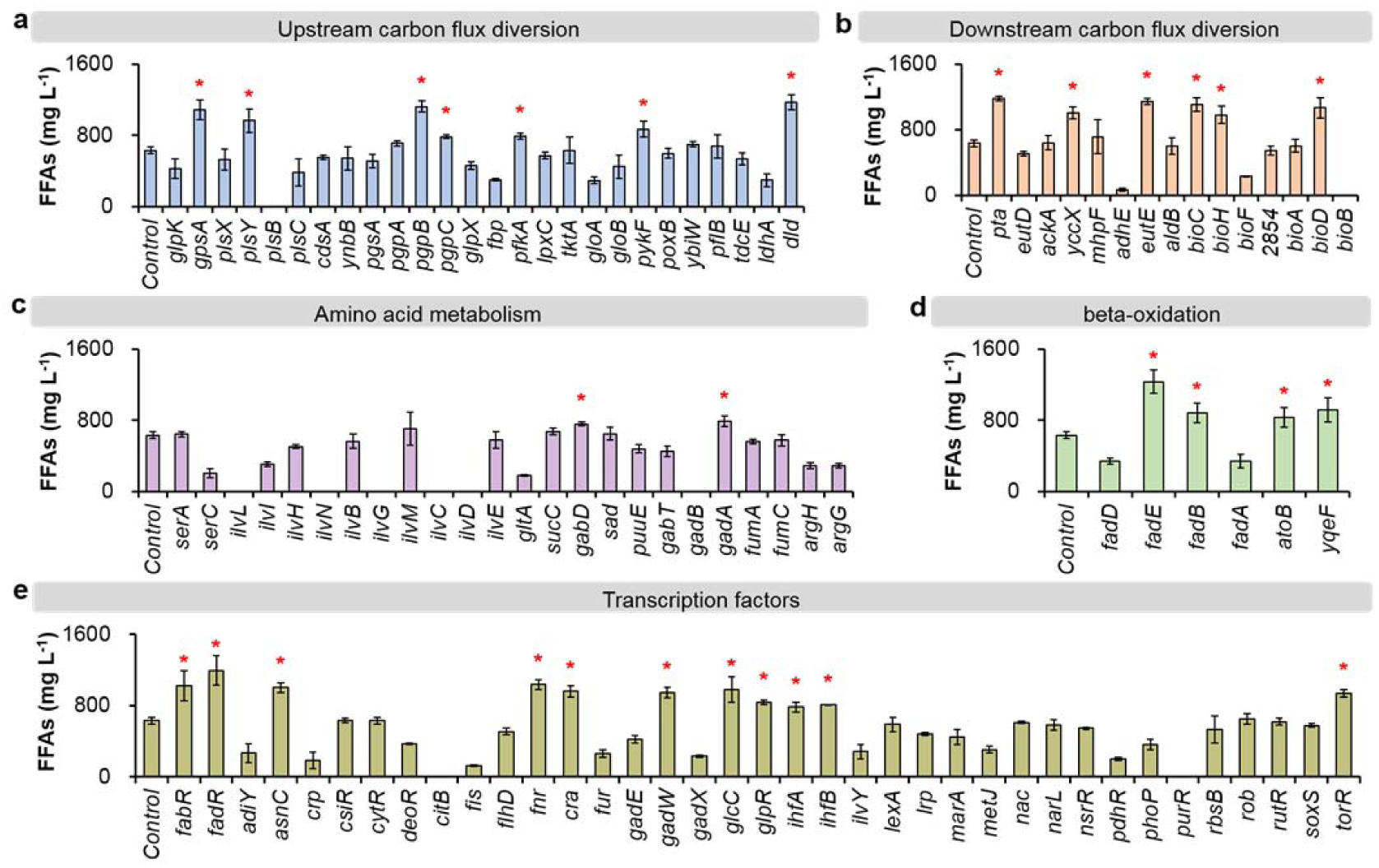
Effect of genetic perturbation by CRISPRi on FFAs production. FFAs production through repression of each gene in upstream carbon flux diversion module (**a**), downstream carbon flux diversion module (**b**), amino acid metabolism module (**c**), beta-oxidation module (**d**), and transcription factors module (**e**) by the CRISPRi system. All target genes were repressed with high efficiency. * represents the strains with more than 20% increase in FFAs production than that of the strain Control (631 mg L^-1^). Error bars indicate the standard deviations of three replicates. Source data are provided as a Source Data file

The identified beneficial genes were almost evenly distributed in the modules of upstream carbon flux diversion (**Fig. 2a**), downstream carbon flux diversion (**Fig. 2b**), beta-oxidation (**Fig. 2d**), and transcription factors (**Fig. 2e**). It suggested that reducing the carbon flux that flows to the by-products phosphatidylglycerol (*gpsA, plsY, pgpB* and *pgpC*), D-Lactate (*dld*), acetate (*pta, yccX* and *eutE*) and biotin (*bioC, bioH* and *bioD*), or impeding fatty acids degradation (*fadE, fadB, atoB* and *yqeF*) contributed to the enhanced production of FFAs. Reducing the expression of a few transcription factors (including *fabR, fadR, asnC, fnr, cra, gadW, glcC, glpR, ihfA, ihfB* and *torR*) also enhanced the production of FFAs (**Fig. 2e**), which might benefit from their global regulation of genes expression. However, reducing amino acids biosynthetic pathways was unfavorable for FFAs production, as the engineered strains obtained decreased titer of FFAs or failed to be constructed (**Fig. 2c**). The possible reason was that these amino acids were necessary for cell viability, but the reduced amount of amino acids upon CRISPRi perturbation was insufficient to maintain normal cell physiology. Thus, in addition to the key enzymes in the FFAs biosynthetic pathway, there exist many chromosomal genes seemingly unrelated to FFAs that could be beneficial to the enhanced FFAs production.

Then, we tuned repression efficiency towards each of the 30 beneficial genes to optimize their individual expression level. sgRNAs were designed to bind the non-template DNA strand at the middle or terminal region to achieve medium or low repression efficiency towards each target^21^ (**Fig. 3a**). The plasmids harboring each of the 60 synthetic sgRNAs were thus individually introduced into the starting strain CF, and the production titer of FFAs was determined, showing that 10 strains (LgpsA, LpfkA, MyccX, LgabD, MgadA, MatoB, LatoB, LglpR, MihfA and LihfA) enabled further improvement in FFAs production (**Fig. 3b**).

**Fig. 3.**
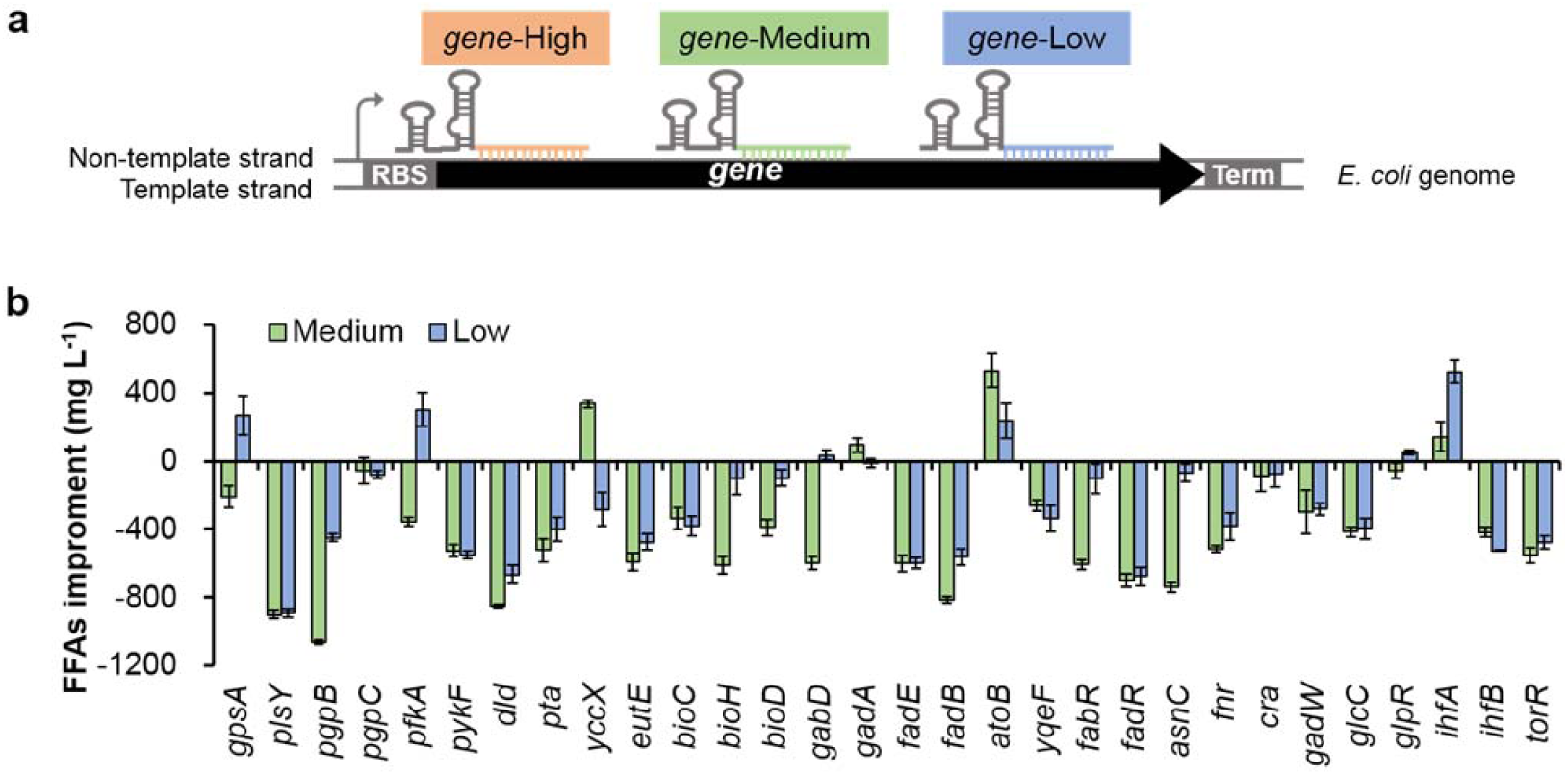
Effect of tuning repression efficiency towards target genes on FFAs production. **a** Design of sgRNA with high, medium or low repression efficiency towards target gene. **b** FFAs production under medium or low repression efficiency compared with that of the high repression efficiency. The repressed genes were screened out from those genes marked with star in Fig. 2. Error bars indicate the standard deviations of three replicates. Source data are provided as a Source Data file

Among the 30 beneficial gene targets, 20 of them (*gpsA, plsY, pgpB, pgpC, dld, yccX, bioC, bioH, bioD, gabD, gadA, asnC, fnr, cra, gadW, glcC, glpR, ihfA, ihfB, torR*) were firstly engineered for FFAs production. We found three favorable metabolic pathways that have not been studied or modulated for FFAs production before, namely the biotin, phosphatidylglycerol, and glutamate biosynthetic pathways. Even in pathways that have been modulated in previous studies, we also identified novel beneficial targets, such as *dld* in the lactate biosynthetic pathway, *yccX* and *eutE* in the acetate formation pathway. In particular, we found a few genes such as *pta* in the acetate formation pathway that have been deleted in previous studies^28^, which however showed no promotional effect on FFAs production, while CRISPRi perturbation of the *pta* expression in this study significantly enhanced FFAs production. Collectively, these results demonstrate that CRISPRi is of use for systematical identification of beneficial targets that can be engineered for improved biosynthesis of a desired product.

### Identification of potential targets in the cellular network via proteome and transcriptome analyses

To explore more potential targets that could facilitate FFAs production, we further focused on transcription factors in consideration of their complicated regulation functions on metabolic networks. Among all the targeted transcription factors, repression of *ihfA* with low efficiency (named as the strain LihfA) achieved a highest FFAs production, followed by the strain HfadR that repressed *fadR* with high efficiency (**Figs. 2e** and **3b**). IhfA is the alpha subunit of the integration host factor (IHF) in *E. coli*, which is capable of affecting transcription on a genome-wide scale^29^. We found that down-regulation of *ihfA* could enhance FFAs production, and the FFAs titer was negatively correlated with the repression efficiency (**Fig. 4a**). Unlike IhfA, FadR controls the expression of several genes directly involved in fatty acid biosynthesis, degradation, and transport across cell membranes^30^. It is intriguing that up-regulation of FadR (named as the strain OfadR) could also increase the FFAs titer as demonstrated in a previous study^31^ and this study (**Fig. 4a**).

**Fig. 4.**
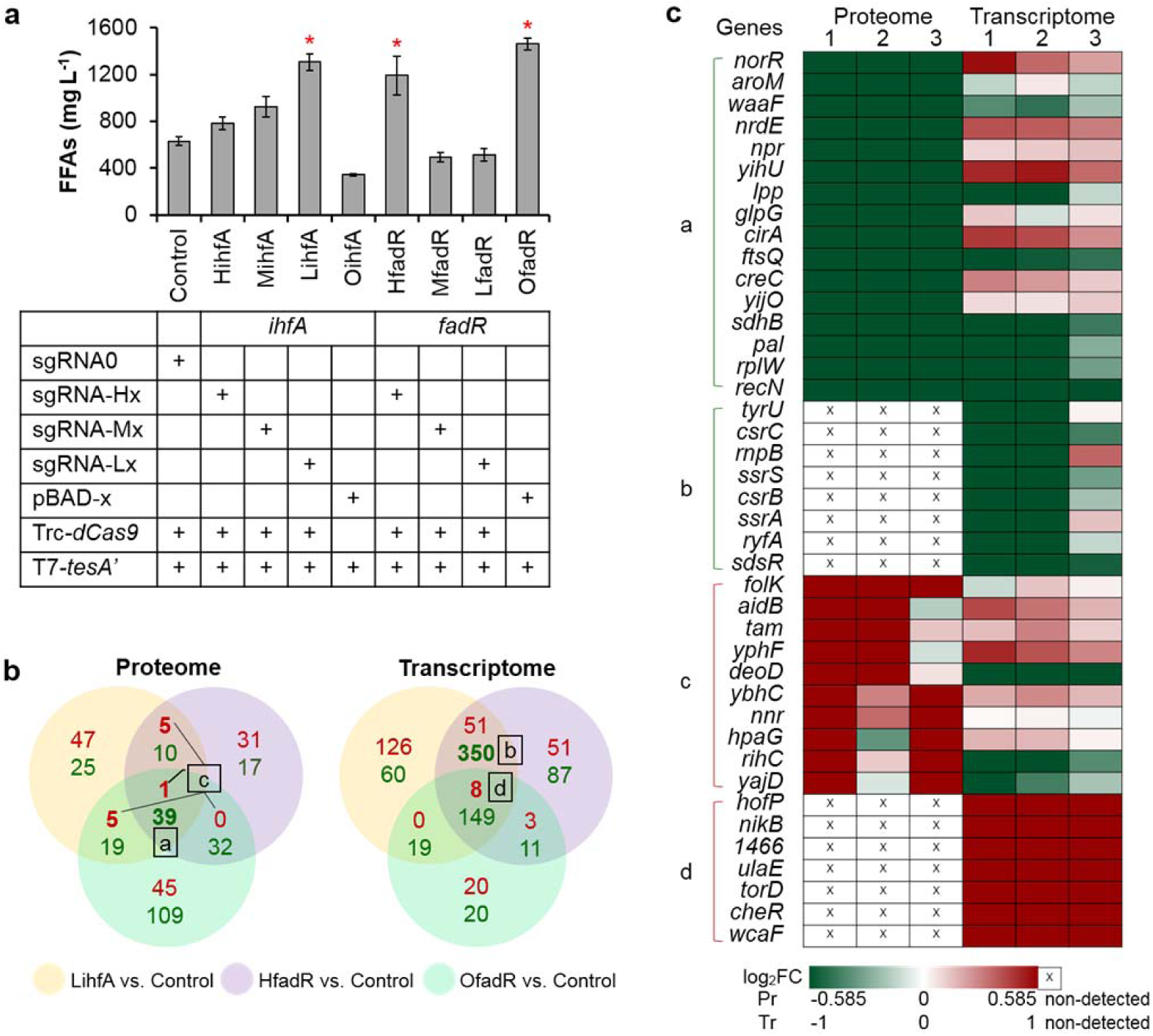
Exploration of potential target genes by proteome and transcriptome analyses. **a** Effect of different *ihfA* or *fadR* expression levels on FFAs production. * represents the strains with high production of FFAs, and these strains were applied to the next omics analyses. **b** A Venn diagram of differentially expressed genes (DEGs). Common DEGs in the high-titer strains at the protein level (a, c) or at the transcript level (b, d) were identified, compared with the strain Control. Green: down-regulated; red: up-regulated. **c** Expression levels of the selected common DEGs in the strains LihfA (1), HfadR (2), and OfadR (3) compared with the strain Control. *1466* represents the gene numbered ECD_01466 in BL21(DE3). Error bars indicate the standard deviations of three replicates. Source data are provided as a Source Data file

Even though the strains with FFAs overproducing phenotype have different genotypes, these strains might have convergent regulatory responses that facilitate FFAs biosynthesis. Thus, the three above engineered strains (LihfA, HfadR and OfadR, with knockdown of *ihfA, fadR* and overexpression of *fadR*, respectively) with high-titer FFAs and the strain Control were applied for proteome and transcriptome analyses to investigate the underlying mechanism of cellular responses to genetic modulations. Cells were sampled at the early stationary phase (24 h upon IPTG induction) (**Supplementary Fig. 3**). Hundreds of genes and proteins were differentially expressed in each pair of LihfA vs. Control, HfadR vs. Control, and OfadR vs. Control (**Supplementary Figs. 4** and **5**), suggesting the global modulation of genes in the engineered strains. To identify convergent rewiring in these high-titer strains, we plotted the differentially expressed genes (DEGs) of the three pairs in a Venn diagram (**Fig. 4b**). As a result, 24 down-regulated targets and 17 up-regulated targets were selected based on their significantly differential expression at the protein level (a and c, respectively) or at the transcript level (b and d, respectively) in at least two pairs (**Fig. 4b**) (see details in Supplementary Materials). The differential expression levels of these 41 genes are summarized in **Fig. 4c**. These potentially favorable gene targets are newly identified and seemingly irrelevant to FFAs biosynthesis in a direct fashion.

To test the function of these 41 candidate targets on FFAs production, the corresponding reverse engineering of each gene was conducted. We applied CRISPRi to repress the expression of the 24 ‘down-regulated’ genes or utilized the P_BAD_ promoter to overexpress the 17 ‘up-regulated’ genes (**Fig. 5a**). The plasmids carrying the cassettes towards each candidate gene were transformed into the starting strain CF. A FFAs assay of the resulting strains showed that modulation of the 19 genes among the identified targets enhanced FFAs production by more than 20% in comparison to the strain Control (**Fig. 5b**). Considering that the selected genes were all differentially expressed in the strain pair of LihfA vs. Control (**Fig. 4c**), these targets were also manipulated in the strain LihfA to test their effects on FFAs production (**Fig. 5c**). As a result, 13 engineered strains gained further increase in FFAs production compared with the strain LihfA (**Fig. 5d**). In particular, the highest FFAs production was obtained in the strain LihfA-OaidB, in which the *ihfA* gene was repressed with low silencing efficiency and the *aidB* gene was overexpressed under the P_BAD_ promoter. The titer is up to 2,052 mg L^-1^, which is 57% and 225% higher than that of the strains LihfA and Control, respectively (**Fig. 5d**). In summary, 26 (*norR, nrdE, npr, yihU, glpG, ftsQ, creC, sdhB, pal, rplW, tyrU, crsC, rnpB, ssrS, csrB, ryfA, sdsR, folk, aidB, yphF, deoD, nnr, rihC, hofP*, ECD_01466, and *ulaE*) out of the 41 newly identified genes are determined as the beneficial targets, which are validated to enhance the FFAs production by reverse engineering.

**Fig. 5.**
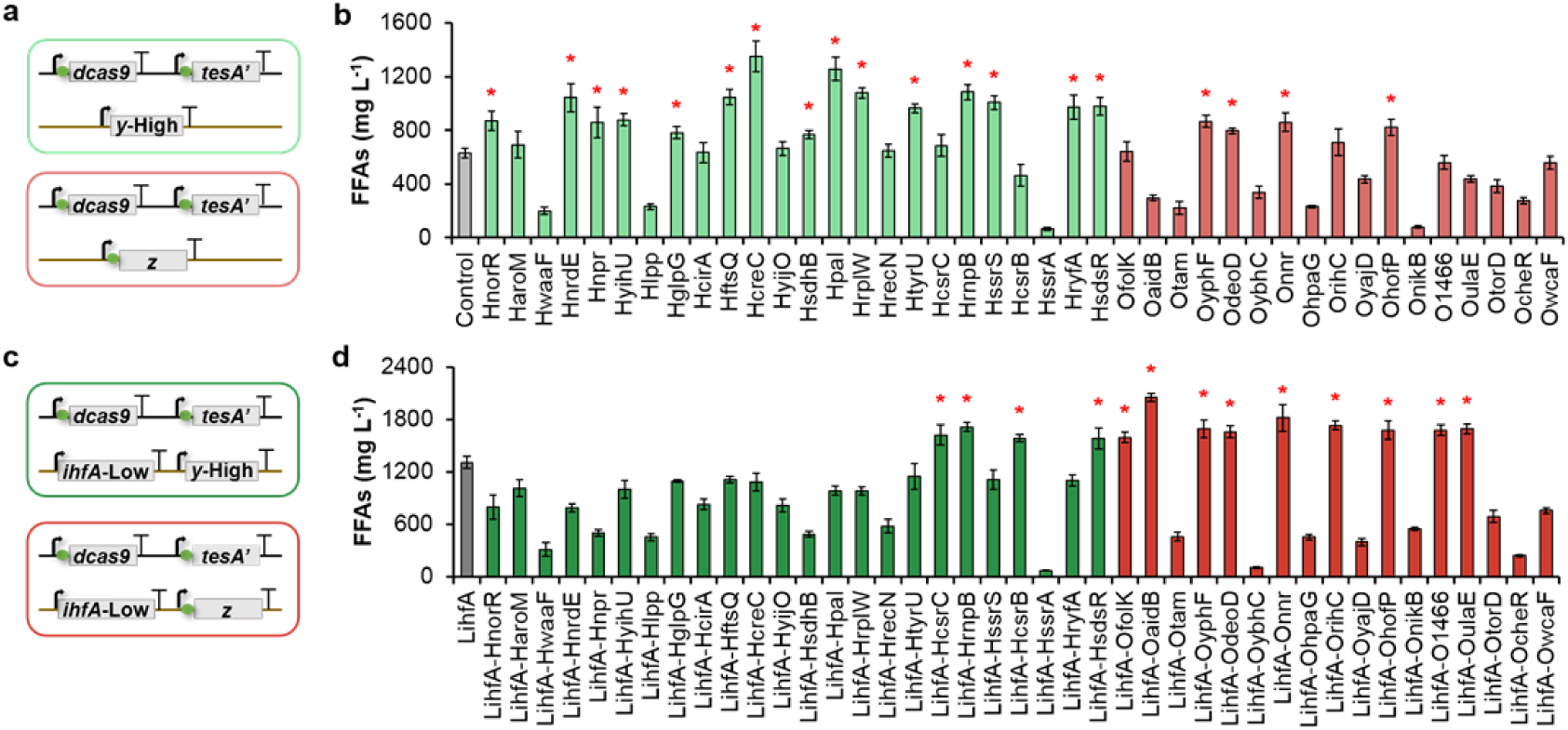
Reverse engineering of the selected DEGs in the strains CF and LihfA. Schematic of cassettes used to repress or overexpress the selected DEGs in the strain CF (**a**), or in the strain LihfA (**c**). The repression was achieved by CRISPRi and the overexpression was enabled by the P_BAD_ promoter. TesA’, dCas9 and sgRNAs were controlled by the P_T7_, P_Trc_ and P_R_ promoter, respectively. *y*: down-regulated gene; *z:* up-regulated gene. Effect of genetic perturbation of the selected DEGs on FFAs production in the strain CF (**b**) or in the strain LihfA (**d**). * represents the strain with FFAs titer increased >20% compared with the reference strain in each bar chart. Green frame or column: repression; red frame or column: overexpression. Error bars indicate the standard deviations of three replicates. Source data are provided as a Source Data file

### Exploration of cellular processes facilitating FFAs production

The newly identified beneficial targets are not in the metabolic pathways that directly related to FFAs biosynthesis. To mine out the related regulatory mechanisms for FFAs production, the functions of the 26 beneficial genes were analyzed according to bioinformatics analysis of the database (NCBI, KEGG, UniProt and Gene Ontology) and summarized into eight sets (**Fig. 6, Supplementary Table 1**). Based on the FFAs titer of the corresponding engineered strains, the weight value (W value) of each set was calculated to characterize their effects on the enhancement of FFAs production. The eight sets were arranged in descending order of W value as follows: cell division, transduction and transport, energy metabolism, cellular structure, non-coding RNAs (ncRNAs), protein metabolism, nucleic acid metabolism, and cofactor metabolism (**Fig. 6, Supplementary Table 1**). Surprisingly, the processes such as cell division, signal transduction, protein metabolism and nucleic acid metabolism seemingly not associated with FFAs biosynthesis in any previous studies that showed favorable effect on FFAs production. These results indicate that the manipulation of cellular processes that are seemingly irrelevant with FFAs biosynthesis can also remarkably boost FFAs production.

**Fig. 6.**
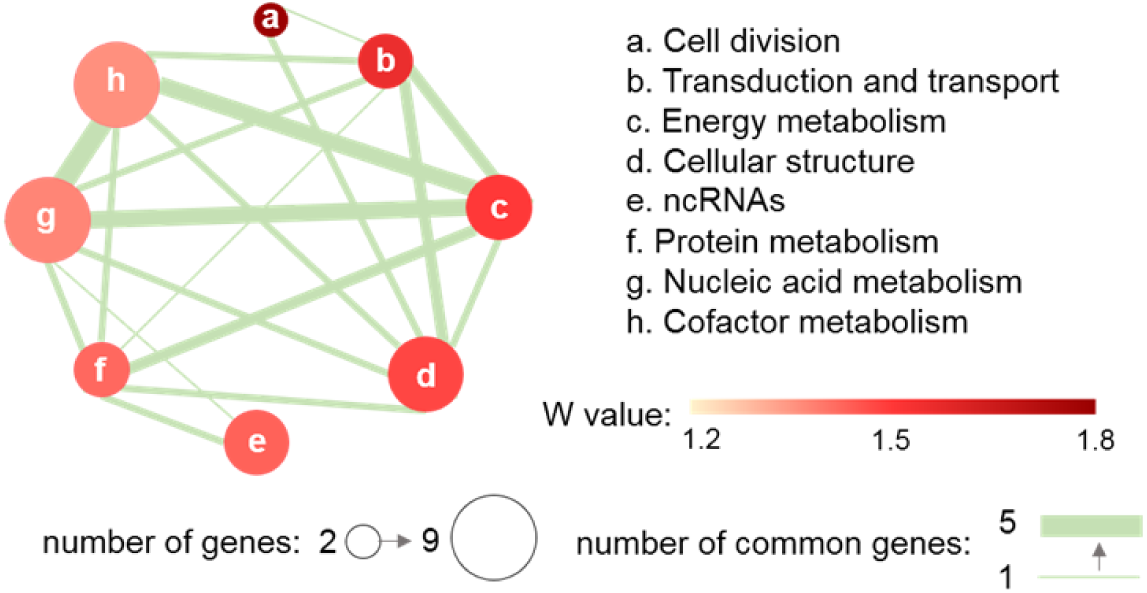
Function network plot of the beneficial genes derived from omics analyses. The size of the node represents the number of genes within the function set. The node’s color represents W value of the function set. The thickness of the edge indicates the number of genes overlapping two function sets. W value is calculated by averaging the FFAs ratio of Hx (Ox) / Control or LiHx (LiOx) / LihfA, x referring to each gene belongs to the function set. Genes and the FFAs ratio are shown in Supplementary Table 1. Source data are provided as a Source Data file

Then, we further dissect the underlying connections between these enriched cellular processes and FFAs production. Down-regulation of *ftsQ* or *pal* significantly increased the FFAs concentration, inactivation of which resulted in deficiency in cell division and formed long multi-septated cell chains^32,33^, which might associated with the increased cell volumes for storing FFAs. Down-regulation of *creC*, a sensor for the early response to phosphate starvation^34^, was favorable for enhanced FFAs production, which is consistent with the promotive effects of phosphate limitation cultivations^35^. The functioning of *npr* down-regulation and *yphF* up-regulation may be associated with the reduction of nitrogen assimilation^36^ and the elevation of carbohydrates’ uptake^37^, indicating a regulatory link between nitrogen and carbon utilization for the efficient FFAs production. Down-regulation of ATP-consuming proteins (such as those encoded by the genes *norR, nrdE, creC* and *rplW*) indirectly increased ATP availability, a potential bottleneck for FFAs biosynthesis^19^. Beneficial effects from down-regulation of membrane proteins (such as those encoded by the genes *glpG, ftsQ, creC* and *pal*) might associated with microbial tolerance to FFAs^38^. The functioning of down-regulation of *rplW* (encoding 50S ribosomal subunit protein L23), *tyrU* (encoding tRNA-Tyr) and *rnpB* (encoding M1 RNA, catalytic component of tRNA processing enzyme RNase P) was associated with the inhibition of protein synthesis, which increased partitioning of total carbon to lipids^8^. Up-regulation of *nnr*, facilitating the repair of NAD(P)H hydrates to NAD(P)H^39^, could be an important complement strategy in enhancing NADPH availability, which plays an important role in FFAs production^19,40^. In all, these findings provide new insights into linkages between cell functions and product biosynthesis, guiding avenues for further strain development.

As a result, the engineered *E. coli* strain LihfA-OaidB with simultaneous repression of *ihfA* and overexpression of *aidB* and *tesA’* achieved the highest titer of FFAs production. According to the results of combined omics analyses and reverse engineering, we concluded that down-regulation of *ihfA* induced comprehensive adjustments of cellular processes that promoted the FFAs production. Nevertheless, repression of *ihfA* might be unfavorable for cellular survival at acidic pH as IHF is required for the induction of acid resistance^41,42^. AidB, one of the DNA damage repairing protein in *E. coli*, is involved in adaptive response of cells, serving to protect cells from cytotoxic effects of acidified cytoplasm or alkylating agents^43-45^. Thus, we speculated that up-regulation of *aidB* in the strain LihfA reconciled the stress from the acidic environment posed by the produced FFAs, facilitating FFAs biosynthesis in the strain LihfA-OaidB.

### Further enhancing FFAs production by fed-batch fermentation

To further enhance FFAs production of the recombinant strain LihfA-OaidB in a scaled-up process, fed-batch fermentations were performed in a 5 L fermenter, using the dO_2_-stat feeding strategy (details described in Methods). With arabinose and IPTG induction, *aidB* (DNA alkylation damage repair protein) was overexpressed by the P_BAD_ promoter, *tesA’* (truncated acyl-CoA thioesterase) was overexpressed by the P_T7_ promoter and *ihfA* (integration host factor subunit alpha) was repressed by CRISPRi with low silencing efficiency in the strain LihfA-OaidB. The FFAs production and cell density of the recombinant strain LihfA-OaidB increased along with the glycerol consumption (**Fig. 7a**). When cells were cultivated for 34 h after IPTG induction, LihfA-OaidB produced a titer of 21.6 g L^-1^ FFAs, accompanied by 0.636 g L^-1^ h^-1^ productivity and 0.146 g FFAs per gram glycerol yield, the highest production titer and productivity in *E. coli* reported to date^18,46^. Extracellularly secreted FFAs were clearly visible at the top of the culture medium after centrifugation (**Fig. 7b**). Moreover, floating dead cells or fatty acid particles^47^ were precipitated and stuck on the fermenter inner wall and sensors over the course of the fermentation process (**Supplementary Fig. 6**), exhibiting an additional production of FFAs that is however hard to accurately quantify and not counted in the FFAs titer. Thus, the overall actual FFAs production was higher than that showed in **Fig. 7a**.

**Fig. 7.**
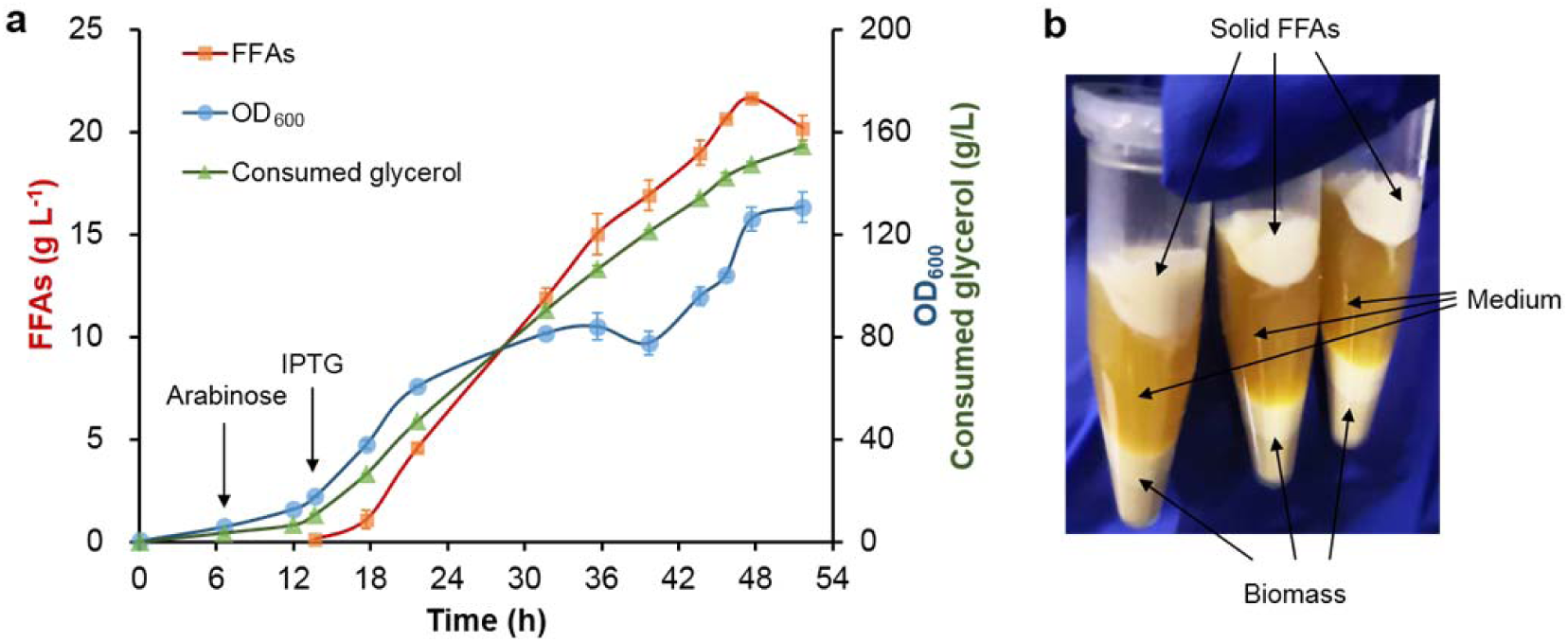
Fed-batch production of FFAs in a 5 L bioreactor. **a** Growth, glucose consumption and FFAs production of the recombinant *E. coli* strain LihfA-OaidB during fed-batch fermentation. **b** Solid FFAs layer after centrifugation. After fermentation, 1 ml of sample was picked up and centrifuged at 5000 g for 10 min. Error bars indicate the standard deviations of three replicates. Source data are provided as a Source Data file

## Discussion

Genome-scale identification of beneficial gene targets in the chromosome for improving production of a desired product is challenging due to the complicated intracellular interactions. We here developed a systems metabolic engineering approach, which took the advantage of CRISPRi to readily and rapidly down-regulate gene expression and the omics analyses to provide potential targets in the cellular interaction network, thus enabling rapid, systematical and effective identification of beneficial targets that can be engineered to optimize FFAs production in *E. coli*. This synergistic strategy integrated the ‘forward’ and ‘reverse’ metabolic engineering cycles and their interaction^48^ (**Supplementary Fig. 7**). In forward metabolic engineering, CRISPRi-mediated gene down-regulation functions were pursued in this work for the identification of beneficial chromosomal targets in the competitive or regulatory pathways of FFAs biosynthesis that can be repressed in high, intermediate and low levels, unlike gene knockout, to enhance FFAs production. In reverse metabolic engineering, proteome and transcriptome analyses of the strains with differential FFAs production provide comprehensive data related to mechanisms of cellular metabolism, which can be validated experimentally via CRISPRi system and plasmid mediated expression system. In doing so, we identified beneficial targets that even seemingly irrelevant with FFAs biosynthesis, which are generally inaccessible to hypothesis-driven or random gene targeting, aiding in expanding hosts’ potential of enhanced products synthesis. To our knowledge, this is the first report that systematically identifies beneficial target genes in the cellular interaction network for enhanced FFAs biosynthesis.

Many new approaches, including in vitro reconstitution of enzymes in the FFAs biosynthetic pathway^49,50^, engineering the reversal of the beta-oxidation pathway^51^, and construction of malonyl-CoA sensors^52,53^, have been studied to enhance FFAs production in *E. coli*. However, these approaches essentially focused on the optimization of FFAs metabolic pathways alone, not exploiting the full potential of interwoven intracellular processes. In our study, gene perturbations on the cellular interaction network, including the competitive pathways of FFAs biosynthesis, transcription factors and cellular processes that seemingly irrelevant with FFAs metabolism, were systematically investigated to unleash the cellular potential and change the cells towards a state of FFAs overproduction. As a result, the best engineered strain LihfA-OaidB produced FFAs with a titer of 21.6 g L^-1^ (0.636 g L^-1^ h^-1^ productivity) under fed-batch conditions, which, to the best of our knowledge, is the maximally reported FFAs titer and productivity by recombinant *E. coli* to date. Furthermore, the potential of the identified beneficial targets and our constructed strains for improving FFAs production could be further integrated and amplified by combinatorial genetic perturbations^54^ and exploiting nongenetic variation^46^.

Beneficial targets that were identified by our synergistic CRISPRi-Omics strategy participate in diverse cellular processes, which support the idea that product synthesis is a complex process and that global optimization of cellular machinery is required for optimized product synthesis. Generally, these targets primarily associate with metabolic processes, other cellular processes or regulatory elements. As to metabolic processes, we identified novel beneficial targets in the common targeted pathways, such as *yccX* and *eutE* in acetate formation pathway, ATP-consuming protein *nrdE* and NAD(P)H-hydrate repair enzyme *nnr*, as well as novel pathways that have not been engineered for FFAs production such as the biotin biosynthetic pathway (*bioC, bioH* and *bioD*). Our target screening approach also emphasizes the importance of other cellular processes or functions in the strain improvement for optimized FFAs production, such as cell division (*ftsQ* and *pal*) related morphology, signaling transduction (*creC*) and transport (*npr* and *yphF*) related resources utilization, cellular structure (*glpG, ftsQ* and *pal*) related cellular tolerance, protein metabolism (*rplW, tyrU, rnpB*) related resources partition, and nucleic acid metabolism (*aidB*) related adaptive response. In addition, we also identified beneficial regulatory elements that have not been engineered for FFAs production, such as the integration host factor *ihfA*, or novel ncRNAs *ryfA* and *sdsR*, which modulate FFAs production through regulation of multiple cellular processes. Thus, our findings provide insights into connections between FFAs biosynthesis and other cellular processes and expand the underlying mechanistic understanding of the regulation of FFAs production in *E. coli*.

Strains obtained through the identification and modulation of genes have proven to be a versatile and robust approach for improved FFAs production, which also offers avenues for engineering more potential targets for further improvement. Knowledge acquired in this work toward understanding the FFAs biosynthetic machinery can be extended to other microorganisms for FFAs production, such as yeast and oleaginous bacteria. The approach for identifying promising modulation candidates is also applicable to other products without obvious screening phenotypes.

In summary, the synergistic CRISPRi-Omics strategy enabling rapid, systematical and effective identification of beneficial targets in the intracellular interaction network provides novel insights and solutions for further increasing productivity of microbial cell factories and better understanding of the link between product biosynthesis and the other processes of cellular machinery.

## Methods

### Experimental materials

Strains and plasmids used in this study are listed in **Supplementary Table 2** and **Supplementary Table 3**, respectively. *E. coli Trans*1-T1 strain was used for cloning. *E. coli* BL21(DE3) and its derived strains were used for fermentation. Plasmids pACYCDuet-1, YX210, YX212 and YX213 were used as the vector for expression of dCas9, GFP or TesA’. Plasmid Sg-S (with two BsaI restriction sites) was constructed and used as a platform vector for expression of various sgRNAs. Plasmid Sg-BAD was used as the vector for expression of chromosome genes. Plasmids, except that for sgRNAs expression, were constructed by standard enzyme digestion and ligation. Target site complementary sequences of sgRNAs were designed according to previous studies^55^. Custom-designed spacers were inserted into plasmid Sg-S by one-step Golden Gate assembly^56,57^, allowing rapid construction of plasmids for expression sgRNAs targeting any genomic locus of interest. Primers annealed to custom-designed spacers and primers used to amplify genes from *E. coli* genomic DNA were listed in **Supplementary Tables 4-7** and **Supplementary Table 8**, respectively. Codon-optimized genes were listed in **Supplementary Table 9**.

### Culturing conditions and mediums

To assess FFAs production by different recombinant *E. coli* strains, fermentation was performed in glass tubes. Each strain was inoculated from a freshly transformed single colony in LB agar plate to 2 mL LB medium as seed culture. When cell accumulation reached to stationary phase, 1% (V/V) of seed culture was re-inoculated to 5 mL modified M9 medium in glass tube. The tube cultures then were induced with 1 mM IPTG when OD_600_ reached ∼1.0 and allowed to grow for an additional 40 h at 30°C and 220 rpm. When the P_BAD_ promoter was applied, 10 mM arabinose was utilized to induce gene expression at OD_600_ 0.5– 0.6. All tube experiments were performed at least in triplicate. Flask fermentation of 50 mL culture broth was implemented for proteome and transcriptome analyses with the identical culturing procedures of tube fermentation. M9 medium used for tube and flask fermentation was as described by Schirmer et al. ^58^ with some modifications: 17.1 g L^-1^ Na_2_HPO_4_·12H_2_O, 3 g L^-1^ KH_2_PO_4_, 0.5 g L^-1^ NaCl, 2 g L^-1^ NH_4_Cl, 2 g L^-1^ yeast extract, 3% glycerol, 0.25 g L^-1^ MgSO_4_·7H_2_O, 11.1 mg L^-1^ CaCl_2_, 10 mg L^-1^ thiamine and 0.1% (v/v) Triton-X100. 1 mL L^-1^ metal trace stock solution was also supplemented, which contained 27 g L^-1^ FeCl_3_·6H_2_O, 2 g L^-1^ ZnCl_2_, 2 g L^-1^ Na_2_MoO_4_·2H_2_O, 1.9 g L^-1^ CuSO_4_·5H_2_O, 0.5 g L^-1^ H_3_BO_3_. pH was adjusted to about 7.2 by Tris. If necessary, 100 μg mL^-1^ carbenicillin or 34 μg mL^-1^ chloramphenicol was supplemented.

For fed-batch fermentation, an overnight LB culture (2 mL) of the freshly transformed single colony was sub-cultured into 200 mL modified mineral medium (6 g L^-1^ NH_4_Cl, 8.5 g L^-1^ KH_2_PO_4_, 0.5 g L^-1^ citrate, 5 g L^-1^ yeast extract, 15 g L^-1^ glycerol, 1 g L^-1^ MgSO_4_·7H_2_O, 0.07 g L^-1^ CaCl_2_·2H_2_O, 4 mL L^-1^ metal trace stock solution, 100 mg L^-1^ thiamine)^59^ with 100 μg mL^-1^ carbenicillin and 34 μg mL^-1^ chloramphenicol. When OD_600_ reached to 3-4, the cultured cells were re-inoculated into a 5 L bioreactor (T&J, China) with 1.8 L modified mineral medium as above. Fermentation temperature was set at 30°C and pH was controlled at 7 by feeding 6 N ammonium hydroxide via an auto pump. Air flow rate was maintained at around 2 L min^-1^. The dissolved oxygen (dO_2_) concentration was controlled above 30% by agitation cascade (300–800 r.p.m). When cell density reached about 6 and 15, 10 mM arabinose and 1 mM IPTG was added into the fermentation cell culture, respectively. Feeding medium (2.47 g L^-1^ MgSO_4_·7H_2_O, 500 g L^-1^ glycerol and 100 g L^-1^ yeast extract) was fed to the fermentation culture after the initial 15 g L^-1^ glycerol was almost depleted and the dissolved oxygen showed a sharp increase. The carbon source restriction strategy was carried out by dO_2_ concentration, given that the dO_2_ level immediately spiked once the carbon source was run out. Once the dO_2_ > 60%, the feeding would automatically start at 5 mL min^-1^ until dO_2_ < 60%. Antifoam (Sigma) was added automatically as needed. Broth samples (∼5 ml) were collected at a series of time points to measure cell density and stored at -20°C for further measurements of residual glycerol and FFAs production.

### GFP Fluorescence assay

Fluorescence signal of GFP was used to characterize repression efficiency of the CRISPRi system. Culturing procedures and induction process of the GFP reporting strains were the same as tube fermentation of the FFAs producing strains. For each tube, 200 μL of sample was diluted into the linear range of the detector with phosphate buffered saline (PBS) after IPTG induction for 20 h. Fluorescence intensity (excitation at 485 nm and emission at 520 nm) and cell density (OD_600_) were detected using 96-well polystyrene plates (black plate with a clear bottom) (Corning Incorporated 3603, USA) and a microplate reader (SpectraMax M2, Molecular Devices, USA). The relative fluorescence intensity was first normalized using OD_600_ and then subtracted that of blank *E. coli* BL21 (DE3). Experiments were performed in triplicate.

### Metabolite extraction and analysis

FFAs titers in whole-cell culture (only FFAs was measured in this study) were quantified following previously published methods^60^. Specifically, 0.5 ml of cell culture (or an appropriate volume of cell culture diluted to 0.5 ml) was acidified with 50 μL of concentrated HCl, spiked with 60 μg of heptadecanoic acid as internal standard. The cell culture was extracted twice with 0.5 ml of ethyl acetate. The extracted FFAs were then determined using a Thermo Scientific TRACE 1300 gas chromatograph (GC) equipped with a TG-WaxMS A column (30 m × 0.32 mm × 0.25 µm; Thermo Scientific) and a Flame Ionization Detector (FID) operating under constant flow rate of the carrier gas (nitrogen) at 1 mL min^-1^. The following temperature program was used: hold at 50°C for 1 min, then heat to 245°C at 30°C min^-1^ and hold at this temperature for 22.5 min. Individual fatty acid species were qualified by authentic homologous standards and quantified by comparing the peak areas with that of the internal standard using the Chromeleon 7.1 software. Total concentrations of free fatty acids were calculated as the sum of C_12_ to C_18_ (saturated and monounsaturated).

Glycerol concentration was determined by high-performance liquid chromatography (HPLC) following Waters standard protocols. Briefly, filtered culture supernatants were analyzed by a Waters HPLC system including a Waters e2695 separation module, a Waters 2414 refractive index detector (RID) and an Aminex HPX-87H column (Bio-Rad). The separation was performed through elution with 5 mM H_2_SO_4_ at a flow rate of 0.6 mL min^-1^ at 65°C for 30 min.

### Proteome analysis

Cells were harvested by centrifugation at 5000 rpm for 5 min at 24 h after IPTG induction and flash frozen with liquid nitrogen. For protein preparation, cells were resuspended in 600 μL lysis buffer (0.05 M Tris, pH 8, 1% SDS, and 8 M Urea) and disrupted by ultrasonication for 5 min (cycles with 2s work and 3s pause). After centrifugation at 13000 rpm and 4°C for 20 min, the supernatant was subjected to protein precipitation by adding pre-cooled acetone with 10 mM DTT. The total precipitate was collected, dried and then re-dissolved in 600 μL Urea buffer (0.05 M Tris, pH 8, and 8 M Urea). The protein samples were quantitated by the Bradford method and qualitated by SDS-PAGE. A total of 100 μg of each sample was trypsin-digested overnight and labeled using 4-plex iTRAQ reagents (AB SCIEX) as manual described. The tryptic peptides corresponding to each sample were labeled with its unique iTRAQ tags and proteomics analysis was conducted using a Triple TOF 6600 system (AB SCIEX). The data were analyzed by Novogene.

### Transcriptome analysis

Cells were harvested after 24 h of IPTG induction by quick centrifugation at 10000 g for 1 min and immediately frozen in liquid nitrogen. Total RNA was extracted using the RNAprep pure Cell/Bacteria Kit (Tiangen) following lysozyme treatment. RNA degradation and contamination were monitored on 1 % agarose gels. RNA concentration was measured by Qubit^®^ RNA Assay Kit in Qubit^®^ 2.0 Flurometer (Life Technologies, CA, USA) and RNA integrity was assessed on a 2100 Bioanalyzer (Agilent Technologies, CA, USA). rRNA is removed using a Ribo-zero kit that leaves the mRNA. A total of 3 μg RNA was used as an input per sample. Sequencing libraries were generated using a NEBNext^®^UltraTM RNA Library Prep Kit for Illumina^®^ (NEB, USA) following the manufacturer’s instructions, and index codes were added to attribute sequences to each sample. Clustering of the index-coded samples was performed on a cBot Cluster Generation System using the TruSeq PE Cluster Kit v3-cBot-HS (Illumina) according to the manufacturer’s instructions. After cluster generation, the library preparations were sequenced on an Illumina Hiseq 4000 platform and paired-end reads were generated. After these steps, the data were analyzed by Novogene.

### Data availability

A reporting summary for this article is available as a Supplementary Information file. The data that support the findings of this study are available from the corresponding author upon request. The source data underlying Figs. 2a-e, 3b, 4a-c, 5b, 5d, 6, and 7a, Supplementary Figs. 1b, 3, 4, and 5 and Supplementary Table 1 are provided as a Source Data file.

## Acknowledgements

This work was supported by the National Key Research and Development Program of China (2018YFA0901300), the National Natural Science Foundation of China (NSFC 21621004), and the Young Elite Scientists Sponsorship Program by Tianjin (TJSQNTJ-2018-16).

## Author contributions

L.F., Y.C. and H.S. conceived and designed the experiments. L.F., J.F., and C.W. performed the plasmids construction experiments. L.F. and J.F. performed the fed-batch fermentation experiments. L.F. and Y.C. analyzed the data. L.F. and Y.C. and H.S. wrote the manuscript. H.S. supervised all aspects of the study.

## Additional information

