## Supplemental Figures and Tables for "Genome-wide targets identification by CRISPRi-Omics for high-titer production of free fatty acids in *Escherichia coli*"

**Select candidate targets from DEGs for reverse engineering.**

As for down-regulated genes, 16 out of the 39 genes (a) were selected based on their significantly down-regulated expression at the protein level. We also picked 8 non-coding RNAs (ncRNAs) out of 350 down-regulated genes at the transcript level (b). As for up-regulated genes, differentially expressed genes at the protein level (c) and at the transcript level (d) were all selected, except *tesA* (which has been engineered in our starting strain) and *ileX* (encoding tRNA-Ile, cannot be regulated by our overexpression cassettes). Hereby we selected 24 down-regulated and 17 up-regulated targets. The differential expression levels of these 41 genes are summarized in Fig. 5c.

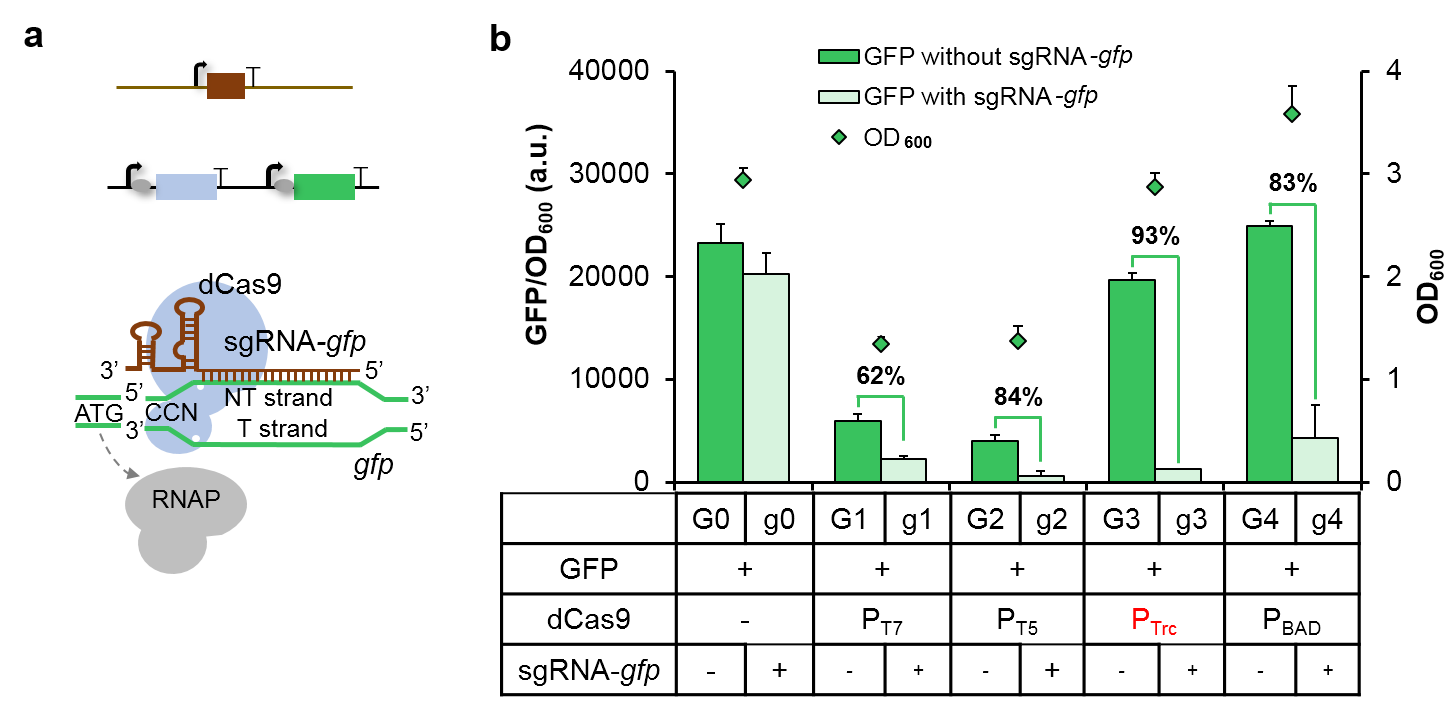

**Supplementary Figure 1. Construction of a non-toxic and functional CRISPRi system in *E. coli* BL21(DE3). a** Schematic of using CRISPRi to block transcription elongation. The cartoon is not to scale. **b** GFP fluorescence-based reporter system was used to test the function of CRISPRi system. dCas9 was expressed under the control of four different promoters. *gfp* and sgRNA-*gfp* were expressed under the control of the P_T7_ promoter and the P_R_ promoter, respectively. Repression efficiency was displayed by numbers above each bar, representing the decreased fluorescence of strain co-expression of *dCas9* and sgRNA-*gfp* (light green bars) compared with that of strain only expressing *dCas9* (dark green bars). Error bars indicate the standard deviations of three replicates. Source data are provided as a Source Data file

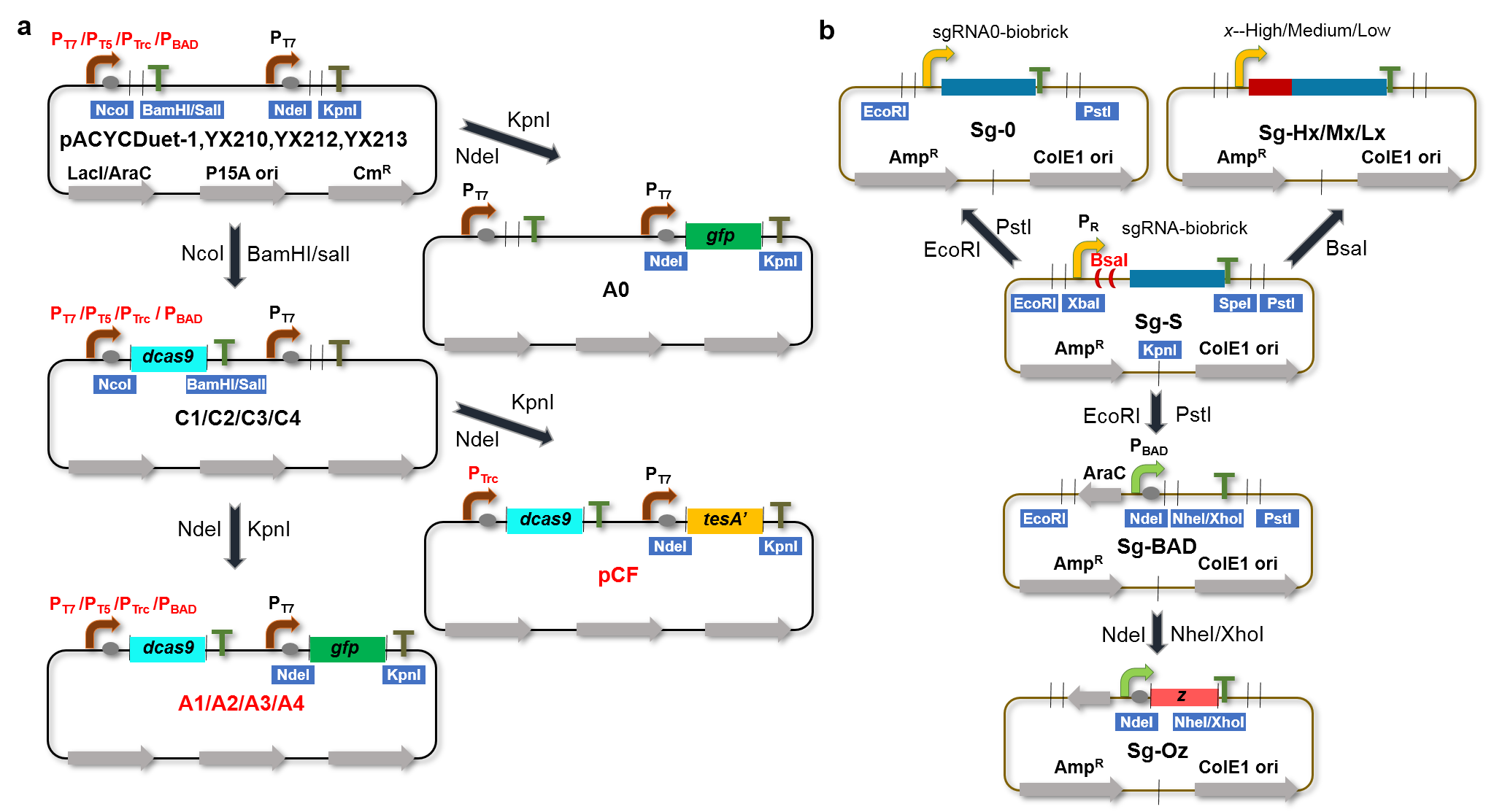

**Supplementary Figure 2. Construction of plasmids. a** Cloning *dcas9* into NcoI and BamHI/SalI site of pACYCDute-1, YX210, YX212 and YX213 resulted in plasmids C1-C4, respectively. And then, subcloning *gfp* into NdeI and KpnI of pACYCDute-1 and plamids C1-C4 resulted in plasmids A0-A4, respectively. Alternatively, subcloning *tesA’* into NdeI and KpnI of plasmid C3 resulted in plasmid pCF. **b** Plasmid Sg-S was constructed via ligation of the synthesized fragments sgRNA-biobrick-ColE1 and Amp-mBsaI after digested by EcoRI and KpnI. Cloning the 20-bp targeting sequence of sgRNA-*gfp* or other sgRNAs into the BsaI of plasmid Sg-S resulted in various sgRNA expression plasmids Sg-Hx/Mx/Lx. Plasmid Sg-0 was constructed via inserting the synthesized fragments sgRNA0-biobrick into plasmid Sg-S by EcoRI and PstI. Alternatively, inserting fragment P_BAD_-T1, amplified from YX213 and digested by EcoRI and PstI, into Sg-S resulted in plasmid Sg-BAD. Genes that need to be upregulated, such as gene *z*, was inserted into Sg-BAD by NdeI and NheI/XhoI, resulting plasmid Sg-Oz.

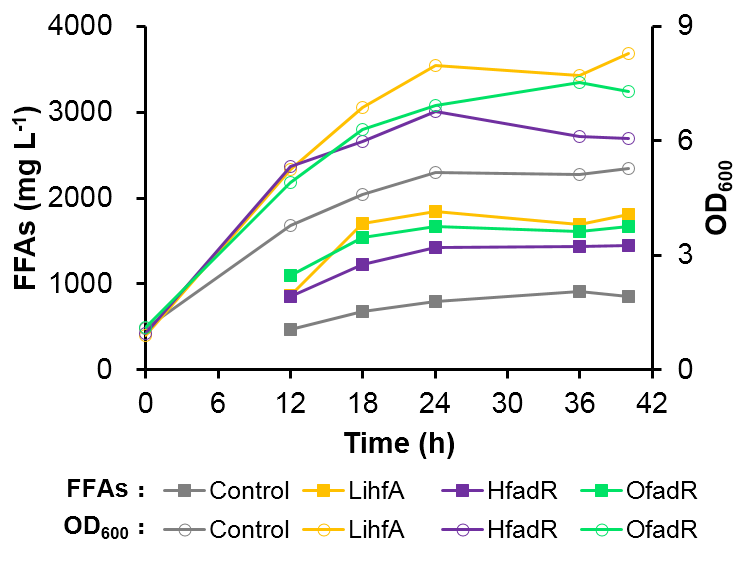

**Supplementary Figure 3.** **Flask shake fermentation of strains sampling for omics analyses.** Time courses of the FFAs titer and cell density (OD_600_) during the fermentation.

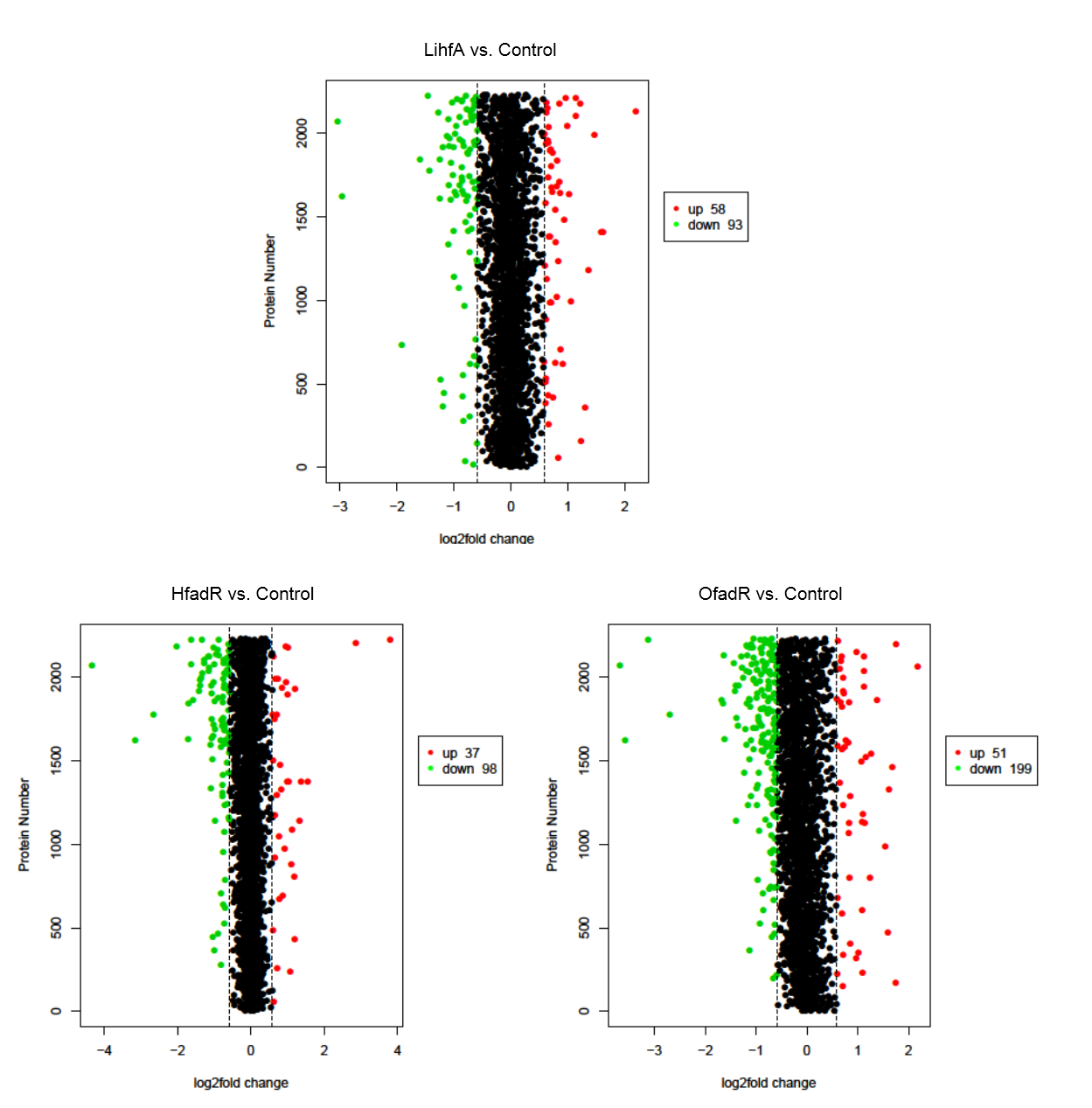

**Supplementary Figure 4. Comparative proteome analysis of the selected strains.** Differentially expressed genes (abs (log_2_ fold change) > 0.585) at the protein level in each strain pair of LihfA vs. Control, HfadR vs. Control and OfadR vs. Control. The red dots mean up-regulated genes and the green dots represent down-regulated genes.

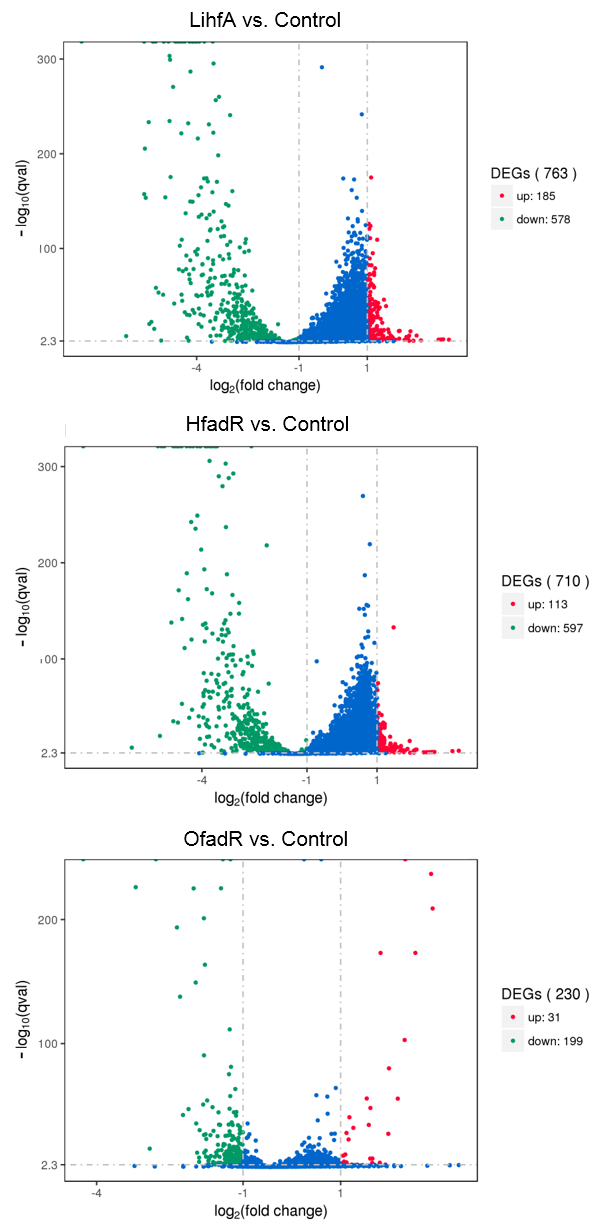

**Supplementary Figure 5. Comparative transcriptome analysis of the selected strains.** Differentially expressed genes at the transcript level (abs (log_2_ fold change) > 1 and q < 0.005) in each strain pair of LihfA vs. Control, HfadR vs. Control and OfadR vs. Control. The red dots mean up-regulated genes and the green dots represent down-regulated genes.

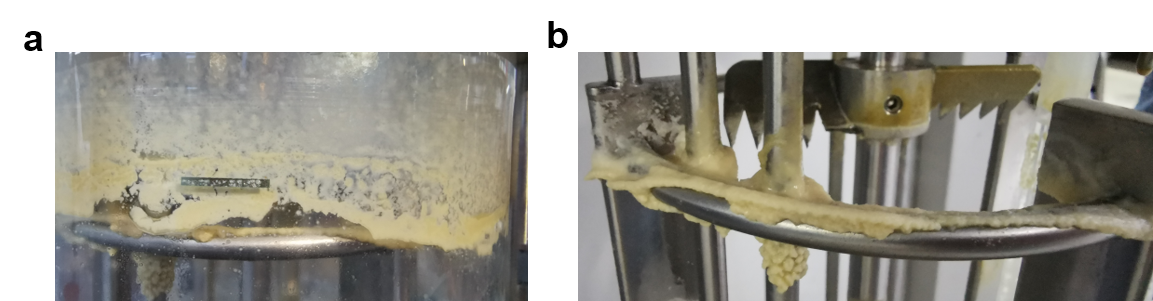

**Supplementary Figure 6. Floating dead cells or fatty acid particles stuck to the fermenter inner wall (a) and sensors (b) during the fermentation.**

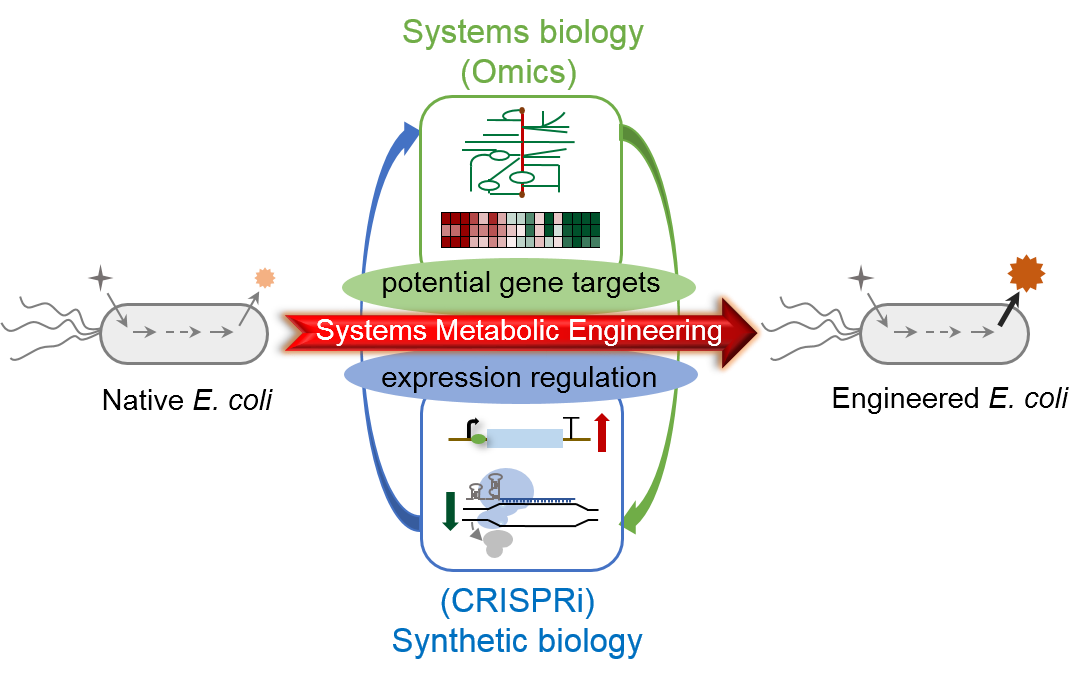

**Supplementary Figure 7. The synergistic CRISPRi-Omics strategy integrated the ‘forward’ and ‘reverse’ metabolic engineering cycles** **and their interaction.** In forward metabolic engineering, CRISPRi-mediated gene down-regulation functions were pursued in this work for the identification of beneficial targets in diverse competitive or regulatory pathways. In reverse metabolic engineering, omics analyses of strains with differential FFAs production provide comprehensive data related to mechanisms of cellular metabolism, which can be validated experimentally via CRISPRi system and plasmid-mediated expression system.

**Supplementary Table 1. Genes in the network and the corresponding FFAs ratio of each engineered strain**

| **Gene** | **FFAs ratio** | | **Function set** |
| --- | --- | --- | --- |
|  | **Hx (Ox)/Control** | **LiHx (LiOx)/LihfA** |  |
| *norR* | 1.3780 |  | bcg |
| *nrdE* | 1.6529 |  | cgh |
| *npr/ptsO* | 1.3556 |  | b |
| *yihU* | 1.3894 |  | bh |
| *glpG* | 1.2390 |  | df |
| *ftsQ* | 1.6577 |  | ad |
| *creC* | 2.1413 |  | bcd |
| *sdhB* | 1.2163 |  | dh |
| *pal* | 1.9911 |  | abd |
| *rplW* | 1.7066 |  | cf |
| *tyrU* | 1.5266 |  | ef |
| *csrC* |  | 1.2400 | e |
| *rnpB* | 1.7175 | 1.3108 | ef |
| *ssrS* | 1.5941 |  | eg |
| *csrB* |  | 1.2125 | e |
| *ryfA* | 1.5396 |  | e |
| *sdsR* | 1.5521 | 1.2102 | e |
| *folK* |  | 1.2186 | cfgh |
| *aidB* |  | 1.5712 | gh |
| *yphF* | 1.3714 | 1.2954 | bcdfgh |
| *deoD* | 1.2595 | 1.2668 | gh |
| *nnr* | 1.3640 | 1.3906 | ch |
| *rihC* |  | 1.3243 | g |
| *hofP* | 1.3034 | 1.2833 | dg |
| *ECD_01466* |  | 1.2836 | d |
| *ulaE* |  | 1.2937 | h |

**^*^**Genes in green are downregulated; genes in red are upregulated; the information sets a to i refer to cell division, transduction and transport, energy metabolism, cellular structure, ncRNAs, protein metabolism, nucleic acid metabolism and cofactor metabolism, respectively.

**Supplementary Table 2. Strains used in this study**

| **Strains** | **Description** | **Reference** |
| --- | --- | --- |
| *E. coli Trans*1-T1 | F^–^ φ80 (*lacZ*) ΔM15 Δ*lacX*74 *hsdR* (*r_k_^–^m_k_^+^*) Δ*recA*1398 *endA*1 *tonA* | TransGen |
| *E. coli* BL21(DE3) | F^–^ *ompT hsdS_B_ (r_B_^–^m_B_^–^) gal dcm* (DE3) | Invitrogen |
| G0 | BL21(DE3) with plasmid A0 | This study |
| G1 | BL21(DE3) with plasmid A1 | This study |
| G2 | BL21(DE3) with plasmid A2 | This study |
| G3 | BL21(DE3) with plasmid A3 | This study |
| G4 | BL21(DE3) with plasmid A4 | This study |
| g0 | BL21(DE3) with plasmids A0 and Sg-gfp | This study |
| g1 | BL21(DE3) with plasmids A1 and Sg-gfp | This study |
| g2 | BL21(DE3) with plasmids A2 and Sg-gfp | This study |
| g3 | BL21(DE3) with plasmids A3 and Sg-gfp | This study |
| g4 | BL21(DE3) with plasmids A4 and Sg-gfp | This study |
| CF (starting strain) | BL21(DE3) with plasmid pCF | This study |
| Control | BL21(DE3) with plasmids pCF and Sg-0 | This study |
| Hx/Mx/Lx | BL21(DE3) with plasmids pCF and Sg-Hx/Mx/Lx | This study |
| Oz | BL21(DE3) with plasmids pCF and Sg-Oz | This study |
| LihfA-Hy | BL21(DE3) with plasmids pCF and Sg-LihfA-Hy | This study |
| LihfA-Oz | BL21(DE3) with plasmids pCF and Sg-LihfA-Oz | This study |

**^*^**x refers to potential knockdown gene targets in metabolic or regulatory pathways related to FFAs biosynthesis. y and z refer to potential knockdown or knockup gene targets identified by omics analyses.

**Supplementary Table 3. Plasmids used in this study**

| **Plasmids** | **Description^*^** | **Reference** |
| --- | --- | --- |
| pACYCDuet-1 | P*_T7_* and P*_T7_*, p15A, Cm^R^ | Novagen |
| YX210 | P*_T5_* and P*_T7_*, p15A, Cm^R^ | Ref. 24 |
| YX212 | P*_Trc_* and P*_T7_*, p15A, Cm^R^ | Ref. 24 |
| YX213 | P*_BAD_* and P*_T7_*, p15A, Cm^R^ | Ref. 24 |
| A0 | P*_T7_*: *gfp*, p15A, Cm^R^ | This study |
| A1 | P*_T7_*: *dcas9*, P*_T7_*: *gfp*, p15A, Cm^R^ | This study |
| A2 | P*_T5_*: *dcas9*, P*_T7_*: *gfp*, p15A, Cm^R^ | This study |
| A3 | P*_Trc_*: *dcas9*, P*_T7_*: *gfp*, p15A, Cm^R^ | This study |
| A4 | P*_BAD_*: *dcas9*, P*_T7_*: *gfp*, p15A, Cm^R^ | This study |
| pCF | P*_Trc_*: *dcas9*, P*_T7_*: *tesA’*, p15A, Cm^R^ | This study |
| Sg-S | P*_R_*: sgRNA cassette, ColE1, Amp^R^ | This study |
| Sg-0 | P*_R_*: sgRNA0 without complementary sequence, ColE1, Amp^R^ | This study |
| Sg-Hx | P*_R_*: sgRNA targets to gene *x* at the front region, ColE1, Amp^R^ | This study |
| Sg-Mx | P*_R_*: sgRNA targets to gene *x* at the middle region, ColE1, Amp^R^ | This study |
| Sg-Lx | P*_R_*: sgRNA targets to gene *x* at the terminal region, ColE1, Amp^R^ | This study |
| Sg-LihfA-Hy | P*_R_*: sgRNA-*ihfA*-Low, P*_R_*: sgRNA-*y*-High, ColE1, Amp^R^ | This study |
| Sg-BAD | P*_BAD_*, ColE1, Amp^R^ | This study |
| Sg-Oz | P*_BAD_*: *z*, ColE1, Amp^R^ | This study |
| Sg-LihfA-Oz | P*_R_*: sgRNA-*ihfA*-Low, P*_BAD_*: *z*, ColE1, Amp^R^ | This study |

^*^Cm^R^, chloramphenicol resistant; Amp^R^, ampicillin resistant.

**Supplementary Table 4. Primers used to construct plasmids Sg-Hx**

| **Primer** | **Sequence (5’→3’)** |
| --- | --- |
| *gfp*-High-F | TTGCcatctaattcaacaagaatt |
| *gfp*-High-R | AAACaattcttgttgaattagatg |
| *glpK*-High-F | TTGCcgcgggagctggtggtgccc |
| *glpK*-High-R | AAACgggcaccaccagctcccgcg |
| *gpsA*-High-F | TTGCcagtcattgaagcattacgt |
| *gpsA*-High-R | AAACacgtaatgcttcaatgactg |
| *plsX*-High-F | TTGCtcccatgacatctaacgcca |
| *plsX*-High-R | AAACtggcgttagatgtcatggga |
| *plsY*-High-F | TTGCagccgcagaggtacgcgatg |
| *plsY*-High-R | AAACcatcgcgtacctctgcggct |
| *plsB*-High-F | TTGCgtagtaaattcgtggccagc |
| *plsB*-High-R | AAACgctggccacgaatttactac |
| *plsC*-High-F | TTGCgactaagatgctgtaaatca |
| *plsC*-High-R | AAACtgatttacagcatcttagtc |
| *cdsA*-High-F | TTGCttaacacaaaagcagatatc |
| *cdsA*-High-R | AAACgatatctgcttttgtgttaa |
| *ynbB*-High-F | TTGCgtaaccgccatagcagaaaa |
| *ynbB*-High-R | AAACttttctgctatggcggttac |
| *pgsA*-High-F | TTGCggaacagtgtaagcaacgta |
| *pgsA*-High-R | AAACtacgttgcttacactgttcc |
| *pgpA*-High-F | TTGCatctttatggcgtggcaaaa |
| *pgpA*-High-R | AAACttttgccacgccataaagat |
| *pgpB*-High-F | TTGCccagaaatccatacggctac |
| *pgpB*-High-R | AAACgtagccgtatggatttctgg |
| *pgpC*-High-F | TTGCcaagtaacgcattcagcggt |
| *pgpC*-High-R | AAACaccgctgaatgcgttacttg |
| *glpX*-High-F | TTGCcgccgctgattcggtgacgc |
| *glpX*-High-R | AAACgcgtcaccgaatcagcggcg |
| *fbp*-High-F | TTGCcagcaaagcagtgagctcac |
| *fbp*-High-R | AAACgtgagctcactgctttgctg |
| *pfkA*-High-F | TTGCacgaacaaccccgcgaattg |
| *pfkA*-High-R | AAACcaattcgcggggttgttcgt |
| *lpxC*-High-F | TTGCcgtcagggtgactttcttgc |
| *lpxC*-High-R | AAACgcaagaaagtcaccctgacg |
| *tktA*-High-F | TTGCccatacccataggggcaccc |
| *tktA*-High-R | AAACgggtgcccctatgggtatgg |
| *gloA*-High-F | TTGCcactttggtataaaaatcga |
| *gloA*-High-R | AAACtcgatttttataccaaagtg |
| *gloB*-High-F | TTGCatgtaattgtcatcaaaggc |
| *gloB*-High-R | AAACgcctttgatgacaattacat |
| *pykF*-High-F | TTGCtccgatggtgcaaacaattt |
| *pykF*-High-R | AAACaaattgtttgcaccatcgga |
| *poxB*-High-F | TTGCcaaaggccgccacttcttcg |
| *poxB*-High-R | AAACcgaagaagtggcggcctttg |
| *ybiW*-High-F | TTGCgagcgtgtccagtttcagtg |
| *ybiW*-High-R | AAACcactgaaactggacacgctc |
| *pflB*-High-F | TTGCggctaacttttcattaagct |
| *pflB*-High-R | AAACagcttaatgaaaagttagcc |
| *tdcE*-High-F | TTGCgtcggcgtacagcttatcgc |
| *tdcE*-High-R | AAACgcgataagctgtacgccgac |
| *ldhA*-High-F | TTGCgtactgttttgtgctataaa |
| *ldhA*-High-R | AAACtttatagcacaaaacagtac |
| *dld*-High-F | TTGCtttattatcagttgttgtca |
| *dld*-High-R | AAACtgacaacaactgataataaa |
| *pta*-High-F | TTGCgaccgacgctggttccggta |
| *pta*-High-R | AAACtaccggaaccagcgtcggtc |
| *ackA*-High-F | TTGCaccatttactgcatcgatga |
| *ackA*-High-R | AAACtcatcgatgcagtaaatggt |
| *eutD*-High-F | TTGCtccggaaaaaccactctggc |
| *eutD*-High-R | AAACgccagagtggtttttccgga |
| *yccX*-High-F | TTGCtcttttcgcttcgtactgtg |
| *yccX*-High-R | AAACcacagtacgaagcgaaaaga |
| *mhpF*-High-F | TTGCaatgttgccagaaccgataa |
| *mhpF*-High-R | AAACttatcggttctggcaacatt |
| *adhE*-High-F | TTGCtacttgctcttgagtgaaac |
| *adhE*-High-R | AAACgtttcactcaagagcaagta |
| *eutE*-High-F | TTGCatctcatgaacggcggcggg |
| *eutE*-High-R | AAACcccgccgccgttcatgagat |
| *aldB*-High-F | TTGCatagcgggtttttaacttaa |
| *aldB*-High-R | AAACttaagttaaaaacccgctat |
| *bioC*-High-F | TTGCtgcatgttgctcatagtgtg |
| *bioC*-High-R | AAACcacactatgagcaacatgca |
| *bioH*-High-F | TTGCatgaacattcccctgacctt |
| *bioH*-High-R | AAACaaggtcaggggaatgttcat |
| *bioF*-High-F | TTGCggcagcacgccgcgcatcga |
| *bioF*-High-R | AAACtcgatgcgcggcgtgctgcc |
| *2854*-High-F | TTGCggaatacacttcatcaatac |
| *2854*-High-R | AAACgtattgatgaagtgtattcc |
| *bioA*-High-F | TTGCatgggtgccagatatggcgt |
| *bioA*-High-R | AAACacgccatatctggcacccat |
| *bioD*-High-F | TTGCccccacttcggtatccgttc |
| *bioD*-High-R | AAACgaacggataccgaagtgggg |
| *bioB*-High-F | TTGCgcgacaatgtccagcgtggg |
| *bioB*-High-R | AAACcccacgctggacattgtcgc |
| *serA*-High-F | TTGCtggtgtaaccagctgcacga |
| *serA*-High-R | AAACtcgtgcagctggttacacca |
| *serC*-High-F | TTGCacctctgccggtagcattgc |
| *serC*-High-R | AAACgcaatgctaccggcagaggt |
| *ilvL*-High-F | TTGCcaggctaatcactcgtagaa |
| *ilvL*-High-R | AAACttctacgagtgattagcctg |
| *ilvI*-High-F | TTGCgcaatgcatcataaatatca |
| *ilvI*-High-R | AAACtgatatttatgatgcattgc |
| *ilvH*-High-F | TTGCcgagtaagactgataatatc |
| *ilvH*-High-R | AAACgatattatcagtcttactcg |
| *ilvN*-High-F | TTGCtacgcccggatggttgcgaa |
| *ilvN*-High-R | AAACttcgcaaccatccgggcgta |
| *ilvB*-High-F | TTGCtcttaatgccctgctgttcc |
| *ilvB*-High-R | AAACggaacagcagggcattaaga |
| *ilvG*-High-F | TTGCgccacccggataaccgaaaa |
| *ilvG*-High-R | AAACttttcggttatccgggtggc |
| *ilvM*-High-F | TTGCaaaacacgttctaaggtttc |
| *ilvM*-High-R | AAACgaaaccttagaacgtgtttt |
| *ilvC*-High-F | TTGCtgcccagctgtgccagctgc |
| *ilvC*-High-R | AAACgcagctggcacagctgggca |
| *ilvD*-High-F | TTGCcatgggtggtggtggcggaa |
| *ilvD*-High-R | AAACttccgccaccaccacccatg |
| *ilvE*-High-F | TTGCaatgtaatcagctttcttcg |
| *ilvE*-High-R | AAACcgaagaaagctgattacatt |
| *gltA*-High-F | TTGCaacagctgtgtccccgttga |
| *gltA*-High-R | AAACtcaacggggacacagctgtt |
| *sucC*-High-F | TTGCgtagtacaggcataacccac |
| *sucC*-High-R | AAACgtgggttatgcctgtactac |
| *gabD*-High-F | TTGCccccgttaatcaacgcctgc |
| *gabD*-High-R | AAACgcaggcgttgattaacgggg |
| *sad*-High-F | TTGCatcgaaattgcatgagttgc |
| *sad*-High-R | AAACgcaactcatgcaatttcgat |
| *puuE*-High-F | TTGCtggcagaaagacgacgctga |
| *puuE*-High-R | AAACtcagcgtcgtctttctgcca |
| *gabT*-High-F | TTGCtgaatttgcccaacaccacg |
| *gabT*-High-R | AAACcgtggtgttgggcaaattca |
| *gadB*-High-F | TTGCacgttttgattctgcgatag |
| *gadB*-High-R | AAACctatcgcagaatcaaaacgt |
| *gadA*-High-F | TTGCggaaatccgttaacagcttc |
| *gadA*-High-R | AAACgaagctgttaacggatttcc |
| *fumA*-High-F | TTGCtcatcttttttgagtggaaa |
| *fumA*-High-R | AAACtttccactcaaaaaagatga |
| *fumC*-High-F | TTGCcgccccacagcttatctgcc |
| *fumC*-High-R | AAACggcagataagctgtggggcg |
| *argH*-High-F | TTGCgaaccgttgatctgctgcct |
| *argH*-High-R | AAACaggcagcagatcaacggttc |
| *argG*-High-F | TTGCtaccaatacgttgacctacc |
| *argG*-High-R | AAACggtaggtcaacgtattggta |
| *fadD*-High-F | TTGCatctccgtcggaacgtccgc |
| *fadD*-High-R | AAACgcggacgttccgacggagat |
| *fadE*-High-F | TTGCgatagaacaacgcgccgagc |
| *fadE*-High-R | AAACgctcggcgcgttgttctatc |
| *fadB*-High-F | TTGCgttgtccagacgaacattaa |
| *fadB*-High-R | AAACttaatgttcgtctggacaac |
| *fadA*-High-F | TTGCtcgccctggcgggtaaccag |
| *fadA*-High-R | AAACctggttacccgccagggcga |
| *atoB*-High-F | TTGCcgcccccaggtcgatggcgc |
| *atoB*-High-R | AAACgcgccatcgacctgggggcg |
| *yqeF*-High-F | TTGCgcaccgcgaaagcagccgat |
| *yqeF*-High-R | AAACatcggctgctttcgcggtgc |
| *fabR*-High-F | TTGCatccttgccaaaagtactgc |
| *fabR*-High-R | AAACgcagtacttttggcaaggat |
| *fadR*-High-F | TTGCactcttccgcgaaacccgcc |
| *fadR*-High-R | AAACggcgggtttcgcggaagagt |
| *adiY*-High-F | TTGCtcagtcaataaaacaataca |
| *adiY*-High-R | AAACtgtattgttttattgactga |
| *asnC*-High-F | TTGCcattgcccattaatgcttcc |
| *asnC*-High-R | AAACggaagcattaatgggcaatg |
| *crp*-High-F | TTGCacaagaaccattcgagagtc |
| *crp*-High-R | AAACgactctcgaatggttcttgt |
| *csiR*-High-F | TTGCctcattcgtaatttttcatc |
| *csiR*-High-R | AAACgatgaaaaattacgaatgag |
| *cytR*-High-F | TTGCcgctttttcaacccgattac |
| *cytR*-High-R | AAACgtaatcgggttgaaaaagcg |
| *deoR*-High-F | TTGCcatctccgaaaccccaagca |
| *deoR*-High-R | AAACtgcttggggtttcggagatg |
| *citB*-High-F | TTGCtcaacgatcaatagggttaa |
| *citB*-High-R | AAACttaaccctattgatcgttga |
| *fis*-High-F | TTGCctgagagttaacggtagaaa |
| *fis*-High-R | AAACtttctaccgttaactctcag |
| *flhD*-High-F | TTGCaatgtgtttcagcaactcgg |
| *flhD*-High-R | AAACccgagttgctgaaacacatt |
| *fnr*-High-F | TTGCgccgtataattcgcttttcc |
| *fnr*-High-R | AAACggaaaagcgaattatacggc |
| *cra*-High-F | TTGCgcccagctgccacggcgttc |
| *cra*-High-R | AAACgaacgccgtggcagctgggc |
| *fur*-High-F | TTGCaacgaggaagcgttactttc |
| *fur*-High-R | AAACgaaagtaacgcttcctcgtt |
| *gadE*-High-F | TTGCtacttttttgatctctgaca |
| *gadE*-High-R | AAACtgtcagagatcaaaaaagta |
| *gadW*-High-F | TTGCtatcgaatgaacgacgaatg |
| *gadW*-High-R | AAACcattcgtcgttcattcgata |
| *gadX*-High-F | TTGCgcgatattcaccattaacca |
| *gadX*-High-R | AAACtggttaatggtgaatatcgc |
| *glcC*-High-F | TTGCctgcaaccacttcgcaaata |
| *glcC*-High-R | AAACtatttgcgaagtggttgcag |
| *glpR*-High-F | TTGCccgccagctcattgaggtcg |
| *glpR*-High-R | AAACcgacctcaatgagctggcgg |
| *ihfA*-High-F | TTGCaccacgcgccgtgctgtaat |
| *ihfA*-High-R | AAACattacagcacggcgcgtggt |
| *ihfB*-High-F | TTGCtctttctatcaattctgact |
| *ihfB*-High-R | AAACagtcagaattgatagaaaga |
| *ilvY*-High-F | TTGCgctttccgccagatgcagga |
| *ilvY*-High-R | AAACtcctgcatctggcggaaagc |
| *lexA*-High-F | TTGCgcgtcggcggcatacctgtc |
| *lexA*-High-R | AAACgacaggtatgccgccgacgc |
| *lrp*-High-F | TTGCtaagaatgttacgatcgata |
| *lrp*-High-R | AAACtatcgatcgtaacattctta |
| *marA*-High-F | TTGCaatagcgtcagtattgcgtc |
| *marA*-High-R | AAACgacgcaatactgacgctatt |
| *metJ-*High-F | TTGCttttaacaccttaagaggaa |
| *metJ*-High-R | AAACttcctcttaaggtgttaaaa |
| *nac*-High-F | TTGCcaatttttacgaagtatttc |
| *nac*-High-R | AAACgaaatacttcgtaaaaattg |
| *narL*-High-F | TTGCtcgggtgatcgtcaatcagc |
| *narL*-High-R | AAACgctgattgacgatcacccga |
| *nsrR*-High-F | TTGCatactggtcatccgcccttc |
| *nsrR*-High-R | AAACgaagggcggatgaccagtat |
| *pdhR*-High-F | TTGCcagttgctgctcaatcacat |
| *pdhR*-High-R | AAACatgtgattgagcagcaactg |
| *phoP*-High-F | TTGCcctgaatctgaactttaagg |
| *phoP*-High-R | AAACccttaaagttcagattcagg |
| *purR*-High-F | TTGCcacgtgtgacacagttgtag |
| *purR*-High-R | AAACctacaactgtgtcacacgtg |
| *rbsB*-High-F | TTGCtagcgcaacagcggaaacca |
| *rbsB*-High-R | AAACtggtttccgctgttgcgcta |
| *rob*-High-F | TTGCtaaaaggtcgcgaataatgc |
| *rob*-High-R | AAACgcattattcgcgacctttta |
| *rutR*-High-F | TTGCatatacagcgcctcttttga |
| *rutR*-High-R | AAACtcaaaagaggcgctgtatat |
| *soxS*-High-F | TTGCctacatcaatgttaagcggc |
| *soxS*-High-R | AAACgccgcttaacattgatgtag |
| *torR*-High-F | TTGCtcaacaataacaatgtgatg |
| *torR*-High-R | AAACcatcacattgttattgttga |

**Supplementary Table 5. Primers used to construct plasmids Sg-Mx**

| **Primer** | **Sequence (5’→3’)** |
| --- | --- |
| *gpsA*-Medium-F | TTGCaatccgggttactgtaaacg |
| *gpsA*-Medium-R | AAACcgtttacagtaacccggatt |
| *plsY*-Medium-F | TTGCcctttaaatccgaagaaaac |
| *plsY*-Medium-R | AAACgttttcttcggatttaaagg |
| *pgpB*-Medium-F | TTGCaaagtgtagaactcatcaac |
| *pgpB*-Medium-R | AAACgttgatgagttctacacttt |
| *pgpC*-Medium-F | TTGCaaccagcggctgcggagagc |
| *pgpC*-Medium-R | AAACgctctccgcagccgctggtt |
| *pfkA*-Medium-F | TTGCaagaagaggtgtcacgcaga |
| *pfkA*-Medium-R | AAACtctgcgtgacacctcttctt |
| *pykF*-Medium-F | TTGCcttgttcgcaaccaaagatc |
| *pykF*-Medium-R | AAACgatctttggttgcgaacaag |
| *dld*-Medium-F | TTGCggtcagcacttccggctggt |
| *dld*-Medium-R | AAACaccagccggaagtgctgacc |
| *pta*-Medium-F | TTGCttggtgttcaccataaatac |
| *pta*-Medium-R | AAACgtatttatggtgaacaccaa |
| *yccX*-Medium-F | TTGCgtcaagatttttggcgtacc |
| *yccX*-Medium-R | AAACggtacgccaaaaatcttgac |
| *eutE*-Medium-F | TTGCttctactaccgcttcgccgc |
| *eutE*-Medium-R | AAACgcggcgaagcggtagtagaa |
| *bioC*-Medium-F | TTGCatacagctcgcggagtgccg |
| *bioC*-Medium-R | AAACcggcactccgcgagctgtat |
| *bioH*-Medium-F | TTGCtcgtcacgagcactaaaaca |
| *bioH*-Medium-R | AAACtgttttagtgctcgtgacga |
| *bioD*-Medium-F | TTGCctcattaccaatgattctat |
| *bioD*-Medium-R | AAACatagaatcattggtaatgag |
| *gabD*-Medium-F | TTGCgcggccaatttcggtcgaac |
| *gabD*-Medium-R | AAACgttcgaccgaaattggccgc |
| *gadA*-Medium-F | TTGCgtgtaggtcacgccgaaagt |
| *gadA*-Medium-R | AAACactttcggcgtgacctacac |
| *fadE*-Medium-F | TTGCcaatagagattttgaactga |
| *fadE*-Medium-R | AAACtcagttcaaaatctctattg |
| *fadB*-Medium-F | TTGCttgatatctttaattctgac |
| *fadB*-Medium-R | AAACgtcagaattaaagatatcaa |
| *atoB*-Medium-F | TTGCttcgagtgacaacatttacc |
| *atoB*-Medium-R | AAACggtaaatgttgtcactcgaa |
| *yqeF*-Medium-F | TTGCgatctcatctttaaatcgtc |
| *yqeF*-Medium-R | AAACgacgatttaaagatgagatc |
| *fabR*-Medium-F | TTGCagcgttcccgcaataataac |
| *fabR*-Medium-R | AAACgttattattgcgggaacgct |
| *fadR*-Medium-F | TTGCcagcacttcctgcgctttat |
| *fadR*-Medium-R | AAACataaagcgcaggaagtgctg |
| *asnC*-Medium-F | TTGCgctttccagctttgccagcg |
| *asnC*-Medium-R | AAACcgctggcaaagctggaaagc |
| *fnr*-Medium-F | TTGCtccagcgcctgagcgaagct |
| *fnr*-Medium-R | AAACagcttcgctcaggcgctgga |
| *cra*-Medium-F | TTGCctgatcggcaccaaccacgc |
| *cra*-Medium-R | AAACgcgtggttggtgccgatcag |
| *gadW*-Medium-F | TTGCcagtatcggtgaatttaccc |
| *gadW*-Medium-R | AAACgggtaaattcaccgatactg |
| *glcC*-Medium-F | TTGCcgagcattttttcataacag |
| *glcC*-Medium-R | AAACctgttatgaaaaaatgctcg |
| *glpR*-Medium-F | TTGCgcggctgcgtaattcgccac |
| *glpR*-Medium-R | AAACgtggcgaattacgcagccgc |
| *ihfA*-Medium-F | TTGCaaacagttcaaccagttctt |
| *ihfA*-Medium-R | AAACaagaactggttgaactgttt |
| *ihfB*-Medium-F | TTGCctcgccctgcgcaagagtcg |
| *ihfB*-Medium-R | AAACcgactcttgcgcagggcgag |
| *torR*-Medium-F | TTGCaccagttcgcgcagttccag |
| *torR*-Medium-R | AAACctggaactgcgcgaactggt |

**Supplementary Table 6. Primers used to construct plasmids Sg-Lx**

| **Primer** | **Sequence (5’→3’)** |
| --- | --- |
| *gpsA*-Low-F | TTGCtaatacttgataaatttcct |
| *gpsA*-Low-R | AAACaggaaatttatcaagtatta |
| *plsY*-Low-F | TTGCaggcaagagagcatcgaaac |
| *plsY*-Low-R | AAACgtttcgatgctctcttgcct |
| *pgpB*-Low-F | TTGCatttcgcgattttcttccgc |
| *pgpB*-Low-R | AAACgcggaagaaaatcgcgaaat |
| *pgpC*-Low-F | TTGCagttgctggagttcaccgcg |
| *pgpC*-Low-R | AAACcgcggtgaactccagcaact |
| *pfkA*-Low-F | TTGCggtgaaccagctgttcgttc |
| *pfkA*-Low-R | AAACgaacgaacagctggttcacc |
| *pykF*-Low-F | TTGCatgagccgttttttcgttgg |
| *pykF*-Low-R | AAACccaacgaaaaaacggctcat |
| *dld*-Low-F | TTGCctgccagtttttccgtttac |
| *dld*-Low-R | AAACgtaaacggaaaaactggcag |
| *pta*-Low-F | TTGCtatcgtcaaccagtgcgcca |
| *pta*-Low-R | AAACtggcgcactggttgacgata |
| *yccX*-Low-F | TTGCaattcccccgagggatgatg |
| *yccX*-Low-R | AAACcatcatccctcgggggaatt |
| *eutE*-Low-F | TTGCctaatacacagcgacgcaga |
| *eutE*-Low-R | AAACtctgcgtcgctgtgtattag |
| *bioC*-Low-F | TTGCggatatcgcccctgctgttg |
| *bioC*-Low-R | AAACcaacagcaggggcgatatcc |
| *bioH*-Low-F | TTGCaagatatatgattcgctgtg |
| *bioH*-Low-R | AAACcacagcgaatcatatatctt |
| *bioD*-Low-F | TTGCggcaaggtttatgtactttc |
| *bioD*-Low-R | AAACgaaagtacataaaccttgcc |
| *gabD*-Low-F | TTGCaccttcacgacccagacccg |
| *gabD*-Low-R | AAACcgggtctgggtcgtgaaggt |
| *gadA*-Low-F | TTGCtgtgtttaaagctgttctgc |
| *gadA*-Low-R | AAACgcagaacagctttaaacaca |
| *fadE*-Low-F | TTGCtcaactttccgcactttctc |
| *fadE*-Low-R | AAACgagaaagtgcggaaagttga |
| *fadB*-Low-F | TTGCaacggtccaccgagaaatgg |
| *fadB*-Low-R | AAACccatttctcggtggaccgtt |
| *atoB*-Low-F | TTGCgaccagaatacgagcaccac |
| *atoB*-Low-R | AAACgtggtgctcgtattctggtc |
| *yqeF*-Low-F | TTGCcaatggtcaatgccacaccc |
| *yqeF*-Low-R | AAACgggtgtggcattgaccattg |
| *fabR*-Low-F | TTGCcacatttcccggaataattg |
| *fabR*-Low-R | AAACcaattattccgggaaatgtg |
| *fadR*-Low-F | TTGCccggcagatttttctgcatc |
| *fadR*-Low-R | AAACgatgcagaaaaatctgccgg |
| *asnC*-Low-F | TTGCggcttgatggtacgcatgat |
| *asnC*-Low-R | AAACatcatgcgtaccatcaagcc |
| *fnr*-Low-F | TTGCtacgcgtatgaccagcaagc |
| *fnr*-Low-R | AAACgcttgctggtcatacgcgta |
| *cra*-Low-F | TTGCagctacggctgagcacgccg |
| *cra*-Low-R | AAACcggcgtgctcagccgtagct |
| *gadW*-Low-F | TTGCgaatgttgcgcaaactgatg |
| *gadW*-Low-R | AAACcatcagtttgcgcaacattc |
| *glcC*-Low-F | TTGCgcaccgccgagcgaatcaaa |
| *glcC*-Low-R | AAACtttgattcgctcggcggtgc |
| *glpR*-Low-F | TTGCacagctccagttgaatatgg |
| *glpR*-Low-R | AAACccatattcaactggagctgt |
| *ihfA*-Low-F | TTGCtgggcgaagcgttttcgacc |
| *ihfA*-Low-R | AAACggtcgaaaacgcttcgccca |
| *ihfB*-Low-F | TTGCcgatcgcgcagttctttacc |
| *ihfB*-Low-R | AAACggtaaagaactgcgcgatcg |
| *torR*-Low-F | TTGCacgacgaattaacacatcga |
| *torR*-Low-R | AAACtcgatgtgttaattcgtcgt |

**Supplementary Table 7. Primers used to construct plasmids Sg-Hy**

| **Primer** | **Sequence (5’→3’)** |
| --- | --- |
| *norR*-High-F | TTGCaatcccacgctgcaattcga |
| *norR*-High-R | AAACtcgaattgcagcgtgggatt |
| *aroM*-High-F | TTGCtaggtacaatgccgatggtc |
| *aroM*-High-R | AAACgaccatcggcattgtaccta |
| *waaF*-High-F | TTGCtcatgtcgccaacccaagac |
| *waaF*-High-R | AAACgtcttgggttggcgacatga |
| *nrdE*-High-F | TTGCttcctgcgtcaggcattctg |
| *nrdE*-High-R | AAACcagaatgcctgacgcaggaa |
| *npr*-High-F | TTGCgatttcaacagtttgcttga |
| *npr*-High-R | AAACtcaagcaaactgttgaaatc |
| *yihU*-High-F | TTGCgataaattcggcatctttag |
| *yihU*-High-R | AAACctaaagatgccgaatttatc |
| *lpp*-High-F | TTGCctgccagcagagtagaaccc |
| *lpp*-High-R | AAACgggttctactctgctggcag |
| *glpG*-High-F | TTGCggttatgttgttgaatcgtg |
| *glpG*-High-R | AAACcacgattcaacaacataacc |
| *cirA*-High-F | TTGCacagcccgacccgtacgaaa |
| *cirA*-High-R | AAACtttcgtacgggtcgggctgt |
| *ftsQ*-High-F | TTGCccagacgcgttccattattg |
| *ftsQ*-High-R | AAACcaataatggaacgcgtctgg |
| *creC*-High-F | TTGCtccgttgctcttcgcacgcc |
| *creC*-High-R | AAACggcgtgcgaagagcaacgga |
| *yijO*-High-F | TTGCaaatctggcgcagggacagc |
| *yijO*-High-R | AAACgctgtccctgcgccagattt |
| *sdhB*-High-F | TTGCagggtgtaatcctgcatacg |
| *sdhB*-High-R | AAACcgtatgcaggattacaccct |
| *pal*-High-F | TTGCcatgccgcaattgccataac |
| *pal*-High-R | AAACgttatggcaattgcggcatg |
| *rplW*-High-F | TTGCtttagcaactttgagtacga |
| *rplW*-High-R | AAACtcgtactcaaagttgctaaa |
| *recN*-High-F | TTGCaacgatagcaaagttgctga |
| *recN*-High-R | AAACtcagcaactttgctatcgtt |
| *tyrU*-High-F | TTGCcgaaccttcgaagtctgtga |
| *tyrU*-High-R | AAACtcacagacttcgaaggttcg |
| *csrC*-High-F | TTGCaatcctctgtcttcgcctcc |
| *csrC*-High-R | AAACggaggcgaagacagaggatt |
| *rnpB*-High-F | TTGCcgaagaggacgacgacgaag |
| *rnpB*-High-R | AAACcttcgtcgtcgtcctcttcg |
| *ssrS*-High-F | TTGCgtggtatgaaatatcggctc |
| *ssrS*-High-R | AAACgagccgatatttcataccac |
| *csrB*-High-F | TTGCgttctcattctccatcctgg |
| *csrB*-High-R | AAACccaggatggagaatgagaac |
| *ssrA*-High-F | TTGCttacgaggccaaccgcccct |
| *ssrA*-High-R | AAACaggggcggttggcctcgtaa |
| *ryfA*-High-F | TTGCcgtttgcgagacggcggaaa |
| *ryfA*-High-R | AAACtttccgccgtctcgcaaacg |
| *sdsR*-High-F | TTGCcatttccctggaccgaatac |
| *sdsR*-High-R | AAACgtattcggtccagggaaatg |

**Supplementary Table 8. Primers used to construct plasmids Sg-Oz**

| **Primer** | **Sequence (5’→3’)^*^** |
| --- | --- |
| NdeI-*ihfA*-F | GAGAGAGCATATGGCGCTTACAAAAGCTGA |
| NheI-*ihfA*-R | ATTGCTAGCTTACTCGTCTTTGGGCGAAG |
| NdeI-*fadR*-F | GAGAGAGCATATGGTCATTAAGGCGCAAAG |
| NheI-*fadR*-R | ATTGCTAGCTTATCGCCCCTGAATGGCTA |
| NdeI-*folk*-F | GACGAAGCATATGACAGTGGCGTATATTG |
| NheI-*folk*-R | GCGGCTAGCTTACCATTTGCTTAATTTGT |
| NdeI-*aidB*-F | CAAGAAGCATATGCACTGGCAAACTCACA |
| NheI-*aidB*-R | ATTGCTAGCTTACACACACACTCCCCCCG |
| NdeI-*tam*-F | GATAGAGCATATGTCTGACTGGAACCCCTC |
| NheI-*tam*-R | ATCGCTAGCTTACTCCATACGCCGGGCAA |
| NdeI-*yphF*-F | GAGAGACCATATGCCTAAAAAAATGAGAAC |
| NheI-*yphF*-R | ATTGCTAGCTTAGGGCAGACCATCAACGT |
| NdeI-*deoD*-F | CACAGAGCATATGGCTACCCCACACATTAA |
| NheI-*deoD*-R | ATTGCTAGCTTACTCTTTATCGCCCAGCAG |
| NdeI-*ybhC*-F | GCCCAGGCATATGAACACATTTTCAGTTTC |
| NheI-*ybhC*-R | ATTGCTAGCTTACTTCTTCGCCTCTGCAA |
| NdeI-*nnr*-F | CAGAGAGCATATGACGGACCATACAATGAA |
| NheI-*nnr*-R | ATTGCTAGCTCAGGGAGCGGGATTACTCG |
| NdeI-*hpaG*-F | GAGAGAGCATATGAAAGGCACTATCTTCG |
| NheI-*hpaG*-R | ATTGCTAGCTCATTTCGCTGTTTCCTCAC |
| NdeI-*rihC*-F | CAGTGACCATATGCGTTTACCTATCTTCCT |
| NheI-*rihC*-R | ATTGCTAGCTTACGGCACCAGAGCCAGC |
| NdeI-*yajD*-F | GAGAGAGCATATGGCTATCATCCCAAAAAA |
| NheI-*yajD*-R | GACGCTAGCTCACTTCTTCTTGTTCATCA |
| NdeI-*hofP*-F | GCTAGAGCATATGAGGGTTAAACGCTGGTT |
| NheI-*hofP*-R | ATTGCTAGCTCATCCACCATCAGCGTCAC |
| NdeI-*nikB*-F | GAGAGAGCATATGTTGCGTTACGTATTAC |
| NheI-*nikB*-R | ATTGCTAGCTCACGCGTGCGCTCCTTCAT |
| NdeI-*1466*-F | GAGAGAGCATATGCATCAGATGCAAACTAC |
| NheI-*1466*-R | ATTGCTAGCTTACTGCATGGTTGCCTTAAT |
| NdeI-*ulaE*-F | GAAGGTCCATATGTTGTCCAAACAAATCCC |
| NheI-*ulaE*-R | ATTGCTAGCTTATGCCGCCTCCACCATGCC |
| NdeI-*torD*-F | TATCCTGCATATGACCACGCTGACAGCACA |
| XhoI-*torD*-R | ATTCTCGAGTTATCTGTTTTGGTGGTCGC |
| NdeI-*cheR*-F | GCATGAGCATATGACTTCATCTCTGCCCTG |
| NheI-*cheR*-R | CAGGCTAGCTTAATCCTTACTTAGCGCAT |
| NdeI-*wcaF*-F | TCGTGCCCATATGCAGGATTTAAGTGGATT |
| NheI-*wcaF*-R | CTTGCTAGCTTATTCAGTTTCAACGCGTTC |

**^*^**Restriction sites are underlined.

**Supplementary Table 9. Sequences of genes**

> *dcas9*	Length: 4115

**ccATGG**ACAAAAAATACTCTATCGGTCTGGCTATCGGTACTAACTCTGTTGGTTGGGCTGTTATCACCGACGAATACAAAGTTCCGTCTAAAAAATTCAAAGTTCTGGGTAACACCGACCGTCACTCTATCAAAAAAAACCTGATCGGTGCTCTGCTGTTCGACTCTGGTGAAACCGCTGAAGCTACCCGTCTGAAACGTACCGCTCGTCGTCGTTACACCCGTCGTAAAAACCGTATCTGCTACCTGCAAGAAATCTTCTCTAACGAAATGGCTAAAGTTGACGACTCTTTCTTCCACCGTCTCGAAGAATCGTTCCTGGTTGAAGAGGACAAAAAACACGAACGTCACCCGATCTTCGGTAACATCGTTGACGAAGTTGCTTACCACGAAAAATACCCGACCATCTACCACCTGCGTAAAAAACTGGTTGACTCTACCGACAAAGCTGACCTGCGTCTGATCTACCTGGCTCTGGCTCACATGATCAAATTCCGTGGTCACTTCCTGATCGAAGGTGACCTGAACCCGGACAACTCTGACGTTGACAAACTGTTCATCCAGCTGGTTCAGACCTACAACCAGCTGTTCGAAGAAAACCCGATCAACGCTTCTGGTGTTGACGCTAAAGCTATCCTGTCTGCTCGTCTGTCTAAATCTCGTCGTCTGGAAAACCTGATCGCTCAGCTGCCGGGTGAAAAAAAAAACGGTCTGTTCGGTAACCTGATCGCTCTGTCTCTGGGTCTGACCCCGAACTTCAAATCTAACTTCGACCTGGCTGAAGACGCTAAACTGCAACTGTCTAAAGACACCTACGACGACGACCTGGACAACCTGCTGGCTCAGATCGGTGACCAGTACGCTGACCTGTTCCTGGCTGCTAAAAACCTGTCTGACGCTATCCTGCTGTCTGACATCCTGCGTGTTAACACCGAAATCACCAAAGCTCCGCTGTCTGCTTCTATGATCAAACGTTACGACGAACACCACCAGGACCTGACCCTGCTGAAAGCTCTGGTTCGTCAGCAGCTGCCGGAAAAATACAAAGAAATCTTCTTCGACCAGTCTAAAAACGGTTACGCTGGTTACATCGACGGTGGTGCTTCTCAGGAAGAATTTTACAAATTCATCAAACCGATCCTGGAAAAAATGGACGGTACTGAAGAACTGCTGGTTAAACTGAACCGTGAAGACCTGCTGCGTAAACAGCGTACCTTCGACAACGGTTCTATCCCGCACCAGATCCACCTGGGTGAACTGCACGCTATCCTGCGTCGTCAGGAAGACTTCTACCCGTTCCTGAAAGACAACCGTGAAAAAATCGAAAAAATCCTGACCTTCCGTATCCCGTACTACGTTGGTCCGCTGGCTCGTGGTAACTCTCGTTTCGCTTGGATGACCCGTAAATCTGAAGAAACCATCACCCCGTGGAACTTCGAAGAAGTTGTTGACAAAGGTGCTTCTGCTCAGTCTTTCATCGAACGTATGACCAACTTCGACAAAAACCTGCCGAACGAAAAAGTTCTGCCGAAACACTCTCTGCTGTACGAATACTTCACCGTTTACAACGAACTGACCAAAGTTAAATACGTTACCGAAGGTATGCGTAAACCGGCTTTCCTGTCTGGTGAACAGAAAAAAGCTATCGTTGACCTGCTGTTCAAAACCAACCGTAAAGTTACCGTTAAACAGCTGAAAGAAGACTACTTCAAAAAAATCGAATGCTTCGACTCTGTTGAAATCTCTGGTGTTGAAGACCGTTTCAACGCTTCTCTGGGTACTTACCACGACCTGCTGAAAATCATCAAAGACAAAGACTTCCTGGACAACGAAGAAAACGAAGACATCCTGGAAGACATCGTTCTGACCCTGACCCTGTTCGAAGACCGTGAAATGATCGAAGAACGTCTGAAAACCTACGCTCACCTGTTCGACGACAAAGTTATGAAACAGCTGAAACGTCGTCGTTACACCGGTTGGGGTCGTCTGTCTCGTAAACTGATCAACGGTATCCGTGACAAACAGTCTGGTAAAACCATCCTGGACTTCCTGAAATCTGACGGTTTCGCTAACCGTAACTTCATGCAGCTGATCCACGACGACTCTCTGACCTTCAAAGAAGACATCCAGAAGGCGCAGGTAAGCGGTCAGGGTGACTCTCTGCACGAACACATCGCTAACCTGGCTGGTTCTCCGGCTATCAAAAAAGGTATCCTGCAAACCGTTAAAGTTGTTGACGAACTGGTTAAAGTTATGGGTCGTCACAAACCGGAAAACATCGTTATCGAAATGGCTCGTGAAAACCAGACCACCCAGAAAGGTCAGAAAAACTCTCGTGAACGTATGAAACGTATCGAAGAAGGTATCAAAGAACTGGGTTCTCAGATCCTGAAAGAACACCCGGTTGAAAACACCCAGCTGCAAAACGAAAAACTGTACCTGTACTACCTGCAAAACGGTCGTGACATGTACGTTGACCAGGAACTGGACATCAACCGTCTGTCTGACTACGACGTTGACGCTATCGTTCCGCAGTCTTTCCTGAAAGACGACTCTATCGACAACAAAGTTCTGACCCGTTCTGACAAAAACCGTGGTAAATCTGACAACGTTCCGTCTGAAGAAGTTGTTAAAAAAATGAAAAACTACTGGCGTCAGCTGCTGAACGCTAAACTGATCACCCAGCGTAAATTCGACAACCTGACCAAAGCTGAACGTGGTGGTCTGTCTGAACTGGACAAAGCTGGTTTCATCAAACGTCAGCTGGTTGAAACCCGTCAGATCACCAAACACGTTGCTCAGATCCTGGACTCTCGTATGAACACCAAATACGACGAAAACGACAAACTGATCCGTGAAGTTAAAGTTATCACCCTGAAATCTAAACTGGTTTCTGACTTCCGTAAAGACTTCCAGTTCTACAAAGTTCGTGAAATCAACAACTACCACCACGCTCACGACGCTTACCTGAACGCTGTTGTTGGTACTGCTCTGATCAAAAAATACCCGAAACTGGAGTCGGAATTTGTGTACGGGGACTACAAAGTTTACGACGTTCGTAAAATGATCGCTAAATCTGAACAGGAAATCGGTAAAGCTACCGCTAAATACTTCTTCTACTCTAACATCATGAACTTCTTCAAAACCGAAATCACCCTGGCTAACGGTGAAATCCGTAAACGTCCGCTGATCGAAACCAACGGTGAAACCGGCGAGATCGTCTGGGACAAAGGCCGTGACTTCGCTACCGTTCGTAAAGTTCTGTCTATGCCGCAGGTTAACATCGTTAAAAAAACCGAAGTTCAGACCGGTGGTTTCTCTAAAGAATCTATCCTGCCGAAACGTAACTCTGACAAACTGATCGCTCGTAAAAAAGACTGGGACCCAAAAAAATACGGTGGCTTCGACTCTCCGACTGTTGCTTACTCTGTTCTGGTTGTTGCTAAAGTTGAAAAAGGTAAATCTAAAAAACTGAAATCTGTTAAAGAACTGCTGGGTATCACCATCATGGAACGTTCTTCTTTCGAAAAAAACCCGATCGACTTCCTGGAAGCTAAAGGTTACAAAGAAGTTAAAAAAGACCTGATCATCAAACTGCCGAAATACTCTCTGTTCGAACTGGAAAACGGTCGTAAACGTATGCTGGCTTCTGCTGGTGAACTGCAAAAAGGTAACGAACTGGCTCTGCCGTCTAAATACGTTAACTTCCTGTACCTGGCTTCTCACTACGAAAAACTGAAAGGTTCTCCGGAAGACAACGAACAGAAACAGCTGTTCGTTGAACAGCACAAACACTACCTGGACGAAATCATCGAACAGATCAGTGAATTTTCTAAACGTGTTATCCTGGCTGACGCTAACCTGGACAAAGTTCTGTCTGCTTACAACAAACACCGTGACAAACCGATCCGTGAACAGGCTGAAAACATCATCCACCTGTTCACCCTGACCAACCTCGGTGCTCCAGCTGCGTTCAAATACTTCGACACCACCATCGACCGTAAACGTTACACCTCTACCAAAGAAGTTCTGGACGCTACCCTGATCCACCAGTCTATCACCGGTCTGTACGAAACCCGTATCGACCTGTCTCAGCTGGGTGGTGACTAA**ggatcc**

> *gfp*	Length: 729

**catATG**CGTAAAGGAGAAGAACTTTTCACTGGAGTTGTCCCAATTCTTGTTGAATTAGATGGTGATGTTAATGGGCACAAATTTTCTGTCAGTGGAGAGGGTGAAGGTGATGCAACATACGGAAAACTTACCCTTAAATTTATTTGCACTACTGGAAAACTACCTGTTCCgTGGCCAACACTTGTCACTACTTTCGGTTATGGTGTTCAATGCTTTGCGAGATACCCAGATCAcATGAAACAGCATGACTTTTTCAAGAGTGCCATGCCCGAAGGTTATGTACAGGAAAGAACTATATTTTTCAAAGATGACGGGAACTACAAGACACGTGCTGAAGTCAAGTTTGAAGGTGATACCCTTGTTAATAGAATCGAGTTAAAAGGTATTGATTTTAAAGAAGATGGAAACATTCTTGGACACAAATTGGAATACAACTATAACTCACACAATGTATACATCATGGCAGACAAACAAAAGAATGGAATCAAAGTTAACTTCAAAATTAGACACAACATTGAAGATGGAAGCGTTCAACTAGCAGACCATTATCAACAAAATACTCCAATTGGCGATGGCCCTGTCCTTTTACCAGACAACCATTACCTGTCCACACAATCTGCCCTTTCGAAAGATCCCAACGAAAAGAGAGACCACATGGTCCTTCTTGAGTTTGTAACAGCTGCTGGGATTACACATGGCATGGATGAACTATACAAATAATAA**ggtacc**

> *tesA’*	Length: 561

**catATG**GCGGACACGTTATTGATTCTGGGTGATAGCCTGAGCGCCGGGTATCGAATGTCTGCCAGCGCGGCCTGGCCTGCCTTGTTGAATGATAAGTGGCAGAGTAAAACGTCGGTAGTTAATGCCAGCATCAGCGGCGACACCTCGCAACAAGGACTGGCGCGCCTTCCGGCTCTGCTGAAACAGCATCAGCCGCGTTGGGTGCTGGTTGAACTGGGCGGCAATGACGGTTTGCGTGGTTTTCAGCCACAGCAAACCGAGCAAACGCTGCGCCAGATTTTGCAGGATGTCAAAGCCGCCAACGCTGAACCATTGTTAATGCAAATACGTCTGCCTGCAAACTATGGTCGCCGTTATAATGAAGCCTTTAGCGCCATTTACCCCAAACTCGCCAAAGAGTTTGATGTTCCGCTGCTGCCCTTTTTTATGGAAGAGGTCTACCTCAAGCCACAATGGATGCAGGATGACGGTATTCATCCCAACCGCGACGCCCAGCCGTTTATTGCCGACTGGATGGCGAAGCAGTTGCAGCCTTTAGTAAATCATGACTCATAA**ggtacc**

> sgRNA-biobrick-ColE1	Length: 1074

**gaattc**GCGGCCGCT**tctaga**GTAACACCGTGCGTGTTGACTATTTTACCTCTGGCGGTGATAATGGTTGCAgagaccAAAggtctcGGTTTTAGAGCTAGAAATAGCAAGTTAAAATAAGGCTAGTCCGTTATCAACTTGAAAAAGTGGCACCGAGTCGGTGCTTTTTTTCAAATAAAACGAAAGGCTCAGTCGAAAGACTGGGCCTTTCGTTTTATCTGTTGTTTGTCGGTGAACT**actagt**AGCGGCCGC**ctgcag**CGTTCGGCTGCGGCGAGCGGTATCAGCTCACTCAAAGGCGGTAATACGGTTATCCACAGAATCAGGGGATAACGCAGGAAAGAACATGTGAGCAAAAGGCCAGCAAAAGGCCAGGAACCGTAAAAAGGCCGCGTTGCTGGCGTTTTTCCATAGGCTCCGCCCCCCTGACGAGCATCACAAAAATCGACGCTCAAGTCAGAGGTGGCGAAACCCGACAGGACTATAAAGATACCAGGCGTTTCCCCCTGGAAGCTCCCTCGTGCGCTCTCCTGTTCCGACCCTGCCGCTTACCGGATACCTGTCCGCCTTTCTCCCTTCGGGAAGCGTGGCGCTTTCTCATAGCTCACGCTGTAGGTATCTCAGTTCGGTGTAGGTCGTTCGCTCCAAGCTGGGCTGTGTGCACGAACCCCCCGTTCAGCCCGACCGCTGCGCCTTATCCGGTAACTATCGTCTTGAGTCCAACCCGGTAAGACACGACTTATCGCCACTGGCAGCAGCCACTGGTAACAGGATTAGCAGAGCGAGGTATGTAGGCGGTGCTACAGAGTTCTTGAAGTGGTGGCCTAACTACGGCTACACTAGAAGGACAGTATTTGGTATCTGCGCTCTGCTGAAGCCAGTTACCTTCGGAAAAAGAGTTGGTAGCTCTTGATCCGGCAAACAAACCACCGCTGGTAGCGGTGGTTTTTTTGTTTGCAAGCAGCAGATTACGCGCAGAAAAAAAGGATCTCAAGAAGATCCTTTGATCTTTTCTACGGGGTCTGACGCTCAGTGGAACGAAAACTCACGTTAAGGGATTTTGGTCATGA**ggtacc**

> Amp-mBsaI	Length: 1154

**ggtacc**GCCTaagcttGCTTGGATTCTCACCAATAAAAAACGCCCGGCGGCAACCGAGCGTTCTGAACAAATCCAGATGGAGTTCTGAGGTCATTACTGGATCTATCAACAGGAGTCCATATATATGAGTAAACTTGGTCTGACAGTTACCAATGCTTAATCAGTGAGGCACCTATCTCAGCGATCTGTCTATTTCGTTCATCCATAGTTGCCTGACTCCCCGTCGTGTAGATAACTACGATACGGGAGGGCTTACCATCTGGCCCCAGTGCTGCAATGATACCGCG**t**GACCCACGCTCACCGGCTCCAGATTTATCAGCAATAAACCAGCCAGCCGGAAGGGCCGAGCGCAGAAGTGGTCCTGCAACTTTATCCGCCTCCATCCAGTCTATTAATTGTTGCCGGGAAGCTAGAGTAAGTAGTTCGCCAGTTAATAGTTTGCGCAACGTTGTTGCCATTGCTACAGGCATCGTGGTGTCACGCTCGTCGTTTGGTATGGCTTCATTCAGCTCCGGTTCCCAACGATCAAGGCGAGTTACATGATCCCCCATGTTGTGCAAAAAAGCGGTTAGCTCCTTCGGTCCTCCGATCGTTGTCAGAAGTAAGTTGGCCGCAGTGTTATCACTCATGGTTATGGCAGCACTGCATAATTCTCTTACTGTCATGCCATCCGTAAGATGCTTTTCTGTGACTGGTGAGTACTCAACCAAGTCATTCTGAGAATAGTGTATGCGGCGACCGAGTTGCTCTTGCCCGGCGTCAATACGGGATAATACCGCGCCACATAGCAGAACTTTAAAAGTGCTCATCATTGGAAAACGTTCTTCGGGGCGAAAACTCTCAAGGATCTTACCGCTGTTGAGATCCAGTTCGATGTAACCCACTCGTGCACCCAACTGATCTTCAGCATCTTTTACTTTCACCAGCGTTTCTGGGTGAGCAAAAACAGGAAGGCAAAATGCCGCAAAAAAGGGAATAAGGGCGACACGGAAATGTTGAATACTCATACTCTTCCTTTTTCAATATTATTGAAGCATTTATCAGGGTTATTGTCTCATGAGCGGATACATATTTGAATGTATTTAGAAAAATAAACAAATAGGGGTTCCGCGCACATTTCCCCGAAAAGTGCCACCTGgagctcGCCT**gaattc**

> sgRNA0-biobrick	Length: 242

**gaattc**GCGGCCGCT**tctaga**GTAACACCGTGCGTGTTGACTATTTTACCTCTGGCGGTGATAATGGTTGCGTTTTAGAGCTAGAAATAGCAAGTTAAAATAAGGCTAGTCCGTTATCAACTTGAAAAAGTGGCACCGAGTCGGTGCTTTTTTTCAAATAAAACGAAAGGCTCAGTCGAAAGACTGGGCCTTTCGTTTTATCTGTTGTTTGTCGGTGAACT**actagt**AGCGGCCGC**ctgcag**

**^*^**Restriction sites are in bold font; sequence of the promoter P_R_ is in orange; sequence of dual BsaI restriction sites is in red; sequence of dCas9 handle and *S. pyogenes* terminator are in blue; sequence of the terminator T1 is in green.
